# Viral communities from long-term anaerobic alkane-oxidizing enrichments may promote cell surface adhesion

**DOI:** 10.1101/2025.07.20.664993

**Authors:** Aditi K. Narayanan, Alon Philosof, Ranjani Murali, Stephanie A. Connon, Gunter Wegener, Victoria J. Orphan

## Abstract

The anaerobic oxidation of methane and higher C2+ alkanes is a dominant metabolism within hydrocarbon-rich deep-sea sediments and is largely mediated by alkane-oxidizing archaea in metabolic partnership with syntrophic sulfate-reducing bacteria. While these processes feed a diverse ecosystem, the viral component of alkane-rich sediments has historically been overlooked. We analyzed the viral community in long-term sediment-free enrichments of alkane-degrading organisms and found that abiotic factors such as incubation temperature had a greater correlation with community composition than with the phylogenetic patterns among individual viral species. No auxiliary metabolic genes directly involved in hydrocarbon oxidation or sulfate reduction were found, but the presence of AMGs involved in heme synthesis pathways common in methane oxidizers hints at a possible viral impact on alkane degradation. We also found evidence supporting the presence of a viral genus infecting several phyla across both the bacterial and archaeal domains, including one of the sulfate-reducing bacterial partners in the alkane-oxidizing syntrophy. Lastly, we report the presence of *nosD*-like proteins in viruses from sediment-derived systems that are not present in water column datasets; their distribution, genomic context, and lack of canonical *nosD* characteristics suggest an alternate adhesion-related role in sediment communities. The number of novel viruses obtained from these enrichment cultures and their potential roles in mediating host physiology illustrate the importance of studying the viral component in laboratory and environmental systems.

## Introduction

Hydrocarbon seepage in marine sediment ecosystems is an important source of energy for chemosynthetic microorganisms and symbiotic animals in the deep sea [1, 2]. As hydrocarbons diffuse or are advected through anoxic sediments and come into contact with sulfate, they are oxidized by a syntrophic partnership of anaerobic archaea and sulfate-reducing bacteria (SRB). The anaerobic oxidation of methane (AOM) occurs in a wide range of environments, including methane cold seeps [3–6], hydrothermally influenced sediments [7–9], terrestrial groundwater [10], mud volcanoes [11], and lake sediments [12–14]. Related archaea performing propane and butane oxidation coupled to sulfate reduction have also been described from hydrothermally-influenced sediments in the Gulf of California [15, 16].

Viruses are an often-overlooked component of seep and vent communities. They are the most abundant biological entities in the ocean, with estimated numbers as high as 10^30^ total viral particles and up to 15 times the number of viral particles as bacteria and archaea [17]. They play a substantial role in community structure, ecology, microbial evolution, and geochemical cycling in the surface ocean and water column [18–23]. It has been estimated that surface and subsurface sediments contain 10-1000 times more viruses than in the pelagic zone on a per volume basis [24]. In low-energy sedimentary environments, viruses are considered the primary agents of microbial mortality [25]. Neglecting the role of viruses in hydrocarbon seeps results in an incomplete picture of marine sediment ecology and biogeochemical processes that influence the fate of methane and higher alkanes in the environment.

Recent work regarding viruses of diverse methane-oxidizing archaea hints at the breadth of potential influence on their ecology and evolution. For example, viral infection is thought to have played a role in the diversification of the ANME-1 clade via the horizontal transfer of the distinct thymidylate synthase *ThyX*, which is non-homologous to the *ThyA* found in other ANME lineages [26]. Other studies in methane seep systems have found viruses with essential metabolic genes, including phosphoadenosine phosphosulfate reductase (*cysH*), which is critical in assimilatory sulfate reduction [27]; O-antigen synthesis, which may be involved in cell-cell adhesion [27]; and several other cofactor-associated genes [28, 29]. Recently, viruses encoding genes exclusive to methane metabolism were found in a variety of methane-rich habitats, including marine and freshwater ecosystems, cow rumen, and permafrost [30]. However, all current viral genomes recovered from hydrocarbon-rich marine sediments have been mined from bulk metagenomes rather than directly sequenced viruses, and few explore the diversity of habitats and metabolic niches in which these organisms are found [31, 32]. Existing work is also limited by the physical complexity of sediment environments and the slow growth of organisms that catalyze the low-energy yielding methane-oxidizing metabolism.

Here, we leverage sediment-free enrichments of organisms from multiple sampling locations at different in-situ temperatures to directly sequence the viral community that is maintained in the presence of these keystone microbial species. These enrichments select for various anaerobic hydrocarbon degraders and their sulfate-reducing partners via the addition of specific substrates. Reducing the complexity of the sediment matrix enables targeted investigations of viruses that persist when hydrocarbon metabolism is dominant. Our findings suggest that temperature and original sampling location influenced community composition far more than they influence the gene content and phylogenetic relationships of individual viruses. Further, we show that while viral genes do not appear to directly influence the processes of hydrocarbon oxidation and sulfate reduction, they may play a role in attachment in biofilm-dominated sediment communities.

## Materials and Methods

A summary of the procedures used is presented here. Detailed methods are available in the supplement under “Supplementary Methods”.

### Development and maintenance of incubations

Anaerobic alkane-oxidizing enrichments from Guaymas Basin, Elba, Menes Caldera, and the Black Sea were developed via dilution of sediment slurries with anoxic artificial seawater medium as described in Reference #[33]. Enrichments from Santa Monica Basin were developed and maintained as described in Reference #[34]. All incubations are maintained in an artificial seawater medium containing sulfate as the sole electron acceptor. See Supplementary Table S1.

### Collection and purification of viruses

Spent media was sampled from enrichment cultures, filtered with 0.22μm filters, concentrated and purified with Optiprep Density Gradient Medium (Cat. D1556, Sigma Aldrich; St. Louis, MO, USA) and ultracentrifugation [35]. VLP counts were determined by SYBR Gold staining and visualization by Fluorescence microscopy.

### Transmission Electron Microscopy

Spent media from incubations were collected, filtered, and concentrated as described under *Collection and purification of viruses*. Five microliters of virus concentrate were spotted on a glow-discharged carbon-coated 300 mesh copper TEM grid, incubated at room temperature for 5 minutes and stained with 2% uranyl acetate before imaging with an FEI Tecnai T12 at 120kV.

### Viral DNA extraction and sequencing

Density-separated fractions were treated with DNaseI to remove free DNA, and viral DNA was extracted as described in the Supplementary Methods. Sequencing libraries were prepared using the Illumina DNA Prep library kit (Cat. 20018705, Illumina, San Diego, CA, USA). The DNA from the two Menes Caldera incubations was pooled to ensure sufficient input for library preparation. Paired-end sequencing was performed at the University of Southern California Keck School of Medicine.

### Cellular DNA extraction and sequencing

Approximately 15mL of each enrichment was collected and centrifuged and the supernatant was removed. The resulting cell pellet was extracted with Qiagen’s DNeasy Blood and Tissue Kit according to their G+ protocol (Cat. #69504, Qiagen, Hilden, Germany). 16S rRNA genes were amplified with Q5 Hot Start Master Mix (Cat. #M0492S, New England Biolabs, Ipswich, MA, USA), using 515F and 926R primers [36] with Illumina adapters for 32 cycles. PCR duplicates were pooled and barcoded with Illumina Nextera XT index2 primers. The sample was sequenced by Laragen (Culver City, CA) using MiSeq Reagent Kit v3 (600-cycle, Cat. #MS-102-3003) on Illumina’s MiSeq platform with the addition of 15-20% PhiX. The 16S rRNA gene sequencing data were not included for Guaymas Basin Methane 1G and Guaymas Basin Methane G due to a significant temporal gap between virus sampling and 16S rRNA gene sampling, in which the bottles were diluted into larger containers. 16S rRNA amplicon data were processed with the DADA2 v.1.22 workflow [37]. Whole-genome library preparation and paired-end sequencing was performed at the Millard and Muriel Jacobs Genetics and Genomics Laboratory, California Institute of Technology.

### Cellular metagenome assembly and binning

Reads were trimmed with bbduk v38.81 [38] and filtered such that reads with <150 nucleotides were removed. Assembly was performed using metaSPAdes version 3.15.3 [39]. Reads were aligned to the assembled contigs using the Burrows-Wheeler Alignment tool version 0.7.17-r1188 [40]before contigs were binned using MetaBAT version 2.12.1 [41] with a minimum contig length of 1500 bp. The CheckM v1.1.3 lineage workflow [42]was used to determine the quality of each bin. Taxonomy was assigned with GTDB-Tk v2.1.0 [43] using the GTDB R207 database [44–47]. All reads were also mapped back to the full set of contigs using bbmap v38.81 [38]; unmapped reads were reassembled with metaSPAdes.

### vMAG assembly, annotation, and phylogeny

Viral reads were trimmed using bbduk v38.81 [38]and contigs were assembled with metaViralSPAdes v3.15.3 [48] as individual samples and as co-assemblies with multiple samples combined. Viral contigs were identified and quality checked using CheckV [49], viralVerify [48], and seeker [50]. Terminase large subunit (*TerL*), Major Capsid Protein (*MCP*) and auxiliary metabolic genes (AMGs) were identified using Vibrant [51]. Relevant alignments were made using MAFFT v7.505 [52], trimmed with trimal v1 4. rev15 [53], and trees were made using IQ-TREE2 [54]. Further details are available in the supplement.

Auxiliary metabolic genes were included in the analysis if classified as such by Vibrant. Because annotations and open reading frame calls are not always reliable without a great deal of manual curation, AMG abundances per sample were quantified as the abundance of the viral contig on which the AMG was found rather than the abundance of the AMG itself. Within a given sample, a contig was included in the AMG calculations if the centered log ratio of the contig’s abundance was >1.

### Viral classification and clustering

Viral contigs that were 98% complete or higher according to CheckV were classified at the family level using GRAViTy [55], with ICTV Virus Metadata Resource VMR_16-180521_MSL36 (https://ictv.global/vmr, [56]). The whole-genome protein clustering network was generated using vConTACT2 [57] using Refseq release 207 and other curated reference phages. vConTACT2 network was visualized using Cytoscape [58].

### Viral abundance calculations and ordination

Viral reads were quantified per viral cluster and per contig using salmon v1.10.0 [59] and summarized using tximport v1.18.0 [60]. DESeq2 v1.30.1 [61] was used to quantify the abundance of each cluster/contig per library. Abundances were centered log ratio (CLR) transformed with ALDEx2 v1.25.1 [62]. Diversity metrics were calculated using the transcripts per million value for each viral cluster, as output by tximport. Principal components, corresponding PERMANOVA (Adonis, analysis of dissimilarities) values, and rank abundances were calculated using the CLR-transformed data. Information on ordination plot generation is available in the Supplementary Methods.

### CRISPR and tRNA identification and host matching

Potential cellular host genomes were obtained from NCBI Projects, metagenomes generated from this study, and personal communications listed in the Extended Methods. Those genomes not generated from this study were chosen because they were sampled from closely-related sites. Sixty-three additional *Archaeoglobus* genomes were obtained from GTDB under the search time “archaeoglobus”.

CRISPRs were detected using CRISPRDetect 3.0 [63] and CRISPRCasTyper v1.1.4 [64]. Host genome CRISPR spacers were combined and dereplicated with MMseqs2 version 13.45111 [65]. BLASTn v2.12.0+ and SpacePharer v5.c2e680a [66] were used to match the viral contigs with host spacers. tRNAs were identified in putative host and viral genomes using tRNAscan-SE v2.0.5 [67] and Aragorn v1.2.41 [68]. Viral and host tRNAs were combined and dereplicated, respectively. Host tRNAs were aligned against viral contigs and viral tRNAs using BLASTn.

Host/phage network diagrams were based on adjacency matrices of viral clusters that shared hosts, and hosts infected by the same viral clusters, respectively. Where viruses were not clustered at the genus level by vConTACT2, the contig was used on its own. Networks were calculated using the networkX from_pandas_adjacency function [69], and self-loops were removed.

### Identification of nosD-like proteins using a hidden Markov model

NosD sequences were identified either based on Vibrant annotation or with the use of a Hidden Markov Model that we generated (procedure detailed in supplement) based on Reference #[70] or from hits from a curated set of genomes. Hits from the viral MAGs in this study, Tara Oceans/Malaspina viral study [71], methane cold seeps [28, 72], lake sediment and water column [73], and soils [74] were retained if they had an hmmsearch e-value<0.001 (HMMER v3.3.2, [75]). Alignments were generated with MUSCLE v3.8.31 [76] and trees were inferred using IQ-TREE [54] using the model finder mode and confidence in tree topology was evaluated with an ultrafast bootstrap value of 1000. Structural models were predicted using Alphafold2 [77] in the monomer mode with reduced databases and visualized using ChimeraX [78–80].

## Results and Discussion

### Low complexity anaerobic alkane-oxidizing enrichment cultures support a rich viral assemblage

Sediment-free enrichment cultures of syntrophic alkane-oxidizing archaea and sulfate-reducing bacteria and an additional enrichment of one of the sulfate-reducing partners (*Desulfofervidus auxilii*, formerly HotSeep-1) were established between 2003 and 2013 at the Max Planck Institute for Marine Microbiology in Bremen, Germany and at the California Institute of Technology in Pasadena, CA. Inocula originated from seep sediments collected at a hydrothermal vent site in Guaymas Basin, Gulf of California in 2009 [7, 15, 81, 82]; Black Sea AOM mats in 2004; a shallow coastal seep off of Elba, Italy in 2010 [83]; the Menes Caldera mud volcano off of Egypt in 2003 [82]; and methane cold seeps in Santa Monica Basin, California in 2013 [34]. All enrichments except for Santa Monica were generated via dilution of bulk seep sediments or, in the case of Elba, coastal sands. Inoculum for the Santa Monica Basin enrichment cultures was created using Percoll density separation and filtration under anoxic conditions to concentrate methane-oxidizing consortia away from sediment particles [34].

Incubations were placed under temperatures similar to those in their native environment and provided with energy substrates intended to enrich hydrocarbon-oxidizing archaea and their sulfate-reducing syntrophic bacterial partners (Figure 1A). The substrates used were hydrogen, methane, propane, and butane, and incubation temperatures spanned from 10°C to 60°C (Supplementary Table S1).

**Figure 1.**
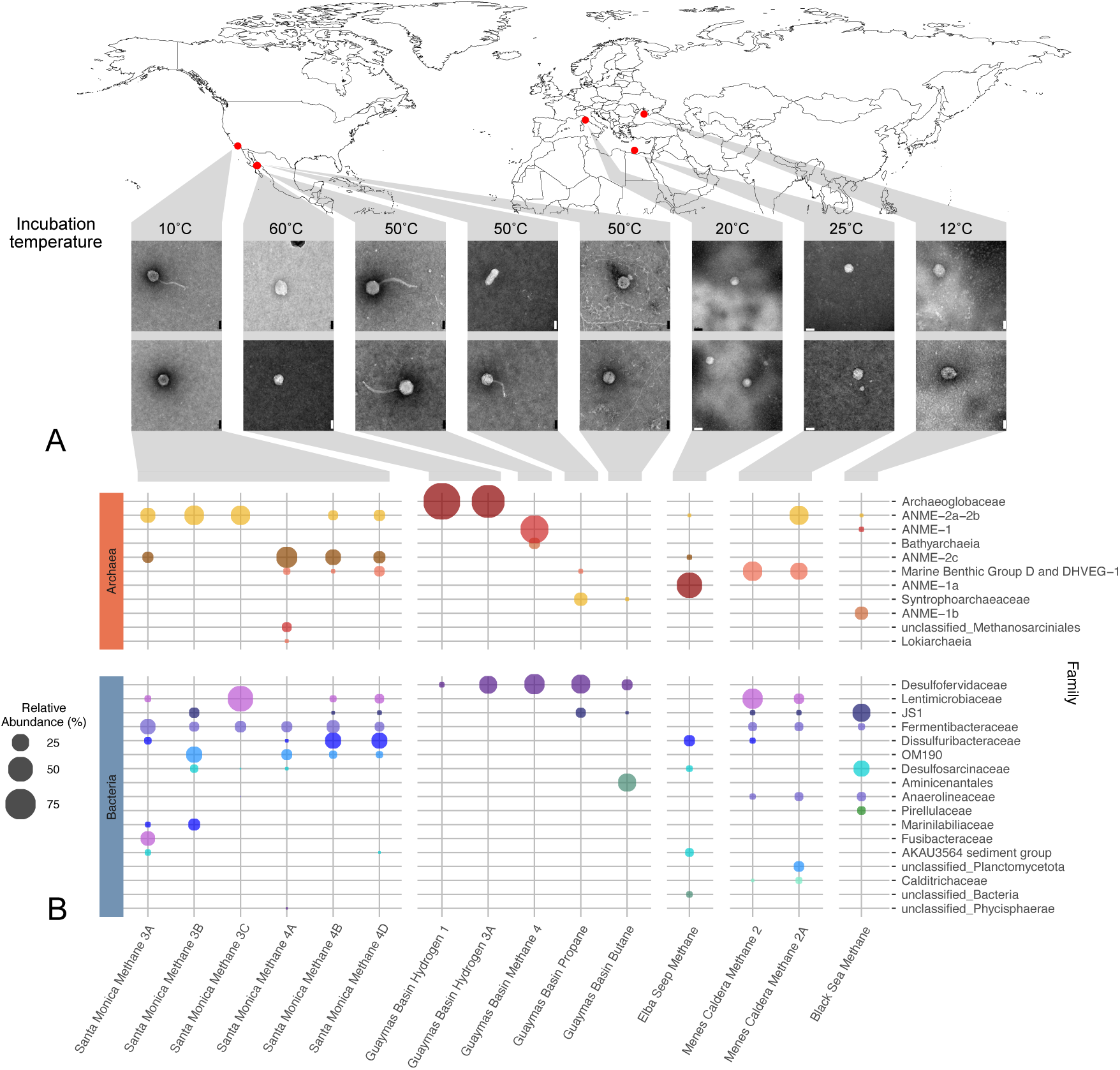
Microbial community composition of enrichment cultures used for metaviromic analysis **A)** Examples of viral-like particles from the enrichment incubations visualized by transmission electron microscopy. Scale bar is 50nm. Corresponding laboratory incubation temperatures are noted and the original sampling location is shown on the map (red symbols). **B)** Family-level archaeal and bacterial taxa detected in enrichment cultures at a relative abundance ≥ 3%, determined by 16S rRNA amplicon sequencing. Samples are grouped by sampling location and were incubated with the listed electron donor. For a complete list of ASVs and their taxonomy, see the Data Availability statement.

16S rRNA amplicon sequencing confirmed the dominant archaeal and bacterial taxa within each incubation bottle (Figure 1B). Major archaeal groups included known methane-oxidizing archaea such as members of the *Methanospirareceae* (ANME-1a,b), *Methanocomedens*, *Methanomarinus*, and *Methanogaster*, (ANME-2a,b,c subgroups); propane/butane-oxidizing *Syntropharchaeaceae*; *Bathyarchaeia* (formerly Miscellaneous Crenarchaeotal Group); *Thermoprofundales* (formerly Marine benthic group D) [84]; *Methanosarcinales,* and *Lokiarchaeota*. The hydrogen amended sulfate-reducing *Desulfofervidus* cultures were unexpectedly overgrown by *Archaeoglobaceae*, an archaeal lineage that was originally reported at low abundance [81]. Major bacterial groups included sulfate-reducing syntrophic bacteria and sulfur disproportionating lineages such as *Desulfofervidaceae* and *Dissulfuribacteraceae*, along with other taxa common in seeps and other anaerobic and/or sedimentary marine environments such as *Fermentibacteraceae*, the JS1/*Atribacteria* group, and *Aminicenantales* [34, 85, 86]. In general, these family-level groups were primarily composed of one or two amplicon sequence variants (ASVs) in each enrichment (Supplementary Table S2).

### Viral ecological patterns differ between the community and OTU levels

Long-term enrichment cultures are valuable systems for investigating key ecological patterns among community members that are difficult to discern directly in the environment, such as viruses. Transmission electron microscopy analysis of concentrated media from all enrichment cultures confirmed the presence of viruses even after several years under controlled laboratory conditions (Figure 1A). We constructed viral genome libraries from concentrated and purified viral-like particles (VLPs; <0.2 µm) from the spent media of the enrichment cultures; in total, 989 dsDNA viral contigs greater than 5000 nucleotides (nt) in length were obtained. Contigs were classified as viral based on identification using ViralVerify [48], Seeker [50], or Vibrant [51], and if more than 50% of the predicted genes were categorized as viral rather than cellular by checkV [49]. Of the 989 viral contigs, 332 were predicted to be at least 98% complete according to checkV. We taxonomically classified these 332 viral contigs using GRAViTy [55] and obtained 60 novel viral families; 110 contigs could not be classified at the family level.

Genus-level clustering of all 989 contigs with vConTACT2 [57] yielded 229 genus-level clusters that contained only 630 contigs, with the remaining 359 contigs unclustered. Despite the reduced complexity of the physicochemical environment (i.e. absence of sediment) and microbial community in our enrichments, a remarkable level of viral richness was maintained (Supplementary Table S3). This pattern aligns with recent findings in an isolate of *E.coli*, in which multiple phages stably coexist in a bacterial culture despite competing for a single host [87]. Nevertheless, the viral diversity recovered across the enrichment cultures was still considerably lower in both taxonomic richness and evenness compared to viral assemblages recovered directly from cold seep sediments [28, 72].

The diversity of origins and incubation conditions of the enrichment cultures provides a unique framework to explore how variables such as temperature, substrate, and original sampling location shape microbial communities. Dimensionality reduction analysis using nonmetric multidimensional scaling (NMDS) of prokaryotic diversity (based on 16S rRNA amplicon sequencing) showed a strong statistical relationship between community composition and factors such as sampling location, incubation temperature, and methane versus non-methane energy sources (Supplementary Table S4). However, the influence of these variables could not åbe disentangled due to their interdependence. A parallel NMDS analysis of the corresponding viral assemblages also showed a significant relationship with both temperature and sampling location (Supplementary Table S4). Interestingly, the source of electron donor (methane vs. non-methane substrates) did not appear to strongly influence the clustering between viral communities (Supplementary Table S4). These patterns remained consistent across different taxonomic resolutions, whether comparing operational taxonomic units (OTUs) or amplicon sequence variants (ASVs) for the prokaryotic communities, or genus-level groupings versus individual contigs for the viruses (Supplementary Figure S1).

To constrain the influence of various physico-chemical parameters on genome-level similarity between viruses, we examined phylogenetic and protein clustering relationships between phage genomes from the different enrichments. Unlike the community-wide PCA analysis shown in Figure 2, phylogenetic analysis of the terminase large subunit (*TerL*) and major capsid protein (*MCP*) showed no discernible relationships between viral clades related to incubation temperature, sampling location, or energy source (Figure 3A, Supplementary Figure S2). As phages lack a universal marker analogous to the 16S rRNA gene, *TerL,* prevalent in tailed viruses, and *MCP* are commonly used proxies [88]. Visualization of vConTACT2 protein clustering networks, generated at the whole-contig level, similarly showed little to no clear pattern (Figure 3B). However, there were differences in how vMAGs clustered between the whole genome network and the *TerL* phylogeny. For example, several groups in the whole-genome network diagram (highlighted in Figure 3B), composed of viruses that share genome-level similarities, were often scattered across the *TerL* phylogeny (corresponding arrows in Figure 3A) with no consistent pattern. Some of these groups identified in the vConTACT2 clustering network showed greater homogeneity with respect to location and incubation temperature, while other groups included a mixture of phage genomes from different sampling sites and incubation temperatures (Figure 3B).

**Figure 2.**
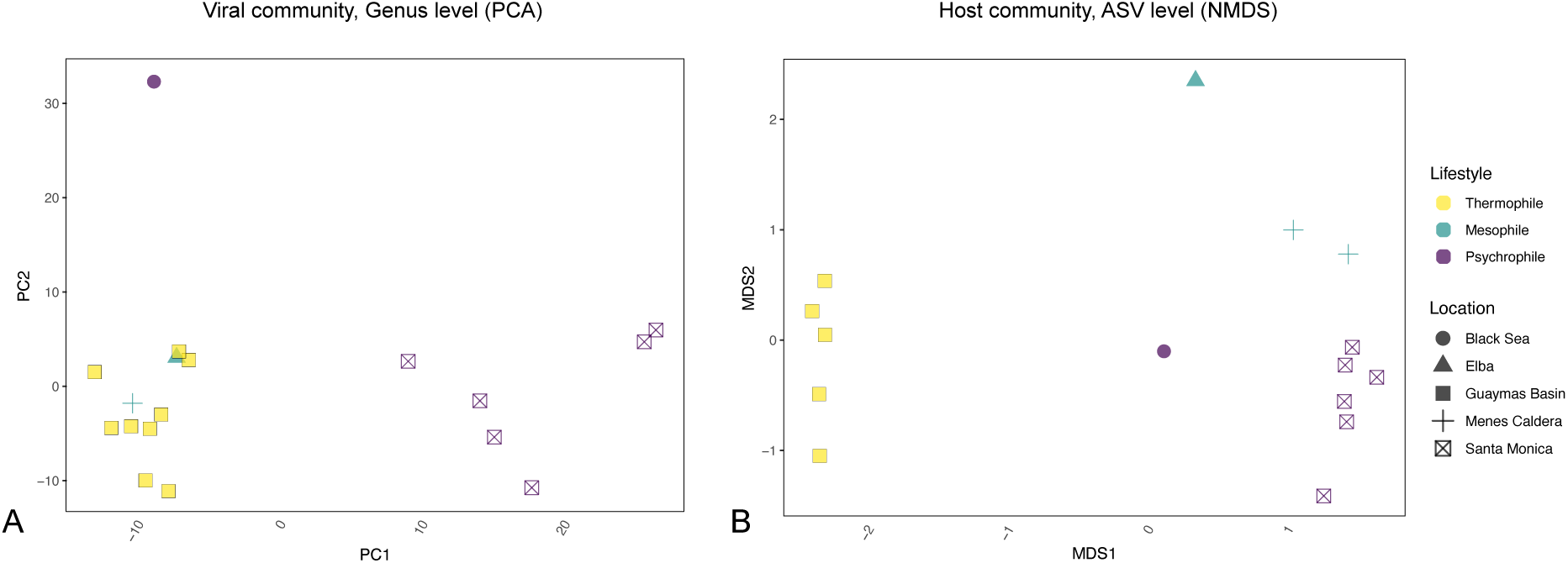
Community-level comparisons between enrichment cultures. **A)** Ordination plot of principal component analysis (PCA) of viral genera recovered from the long-term alkane-oxidizing enrichment cultures. PC1 importance: 0.3349. PC2 importance: 0.1638. **B)** Ordination plot of nonmetric multidimensional scaling (NMDS) of co-occurring microbial 16S rRNA genes at ASV level. Stress: 0.065. Colours represent temperature-based designation for the enriched community and the different symbols indicate the original site location. Note that for the viral community, DNA from the two Menes Caldera incubations was combined for the sequencing library (see Methods), resulting in one data point for the viral community though there are two for the host.

**Figure 3.**
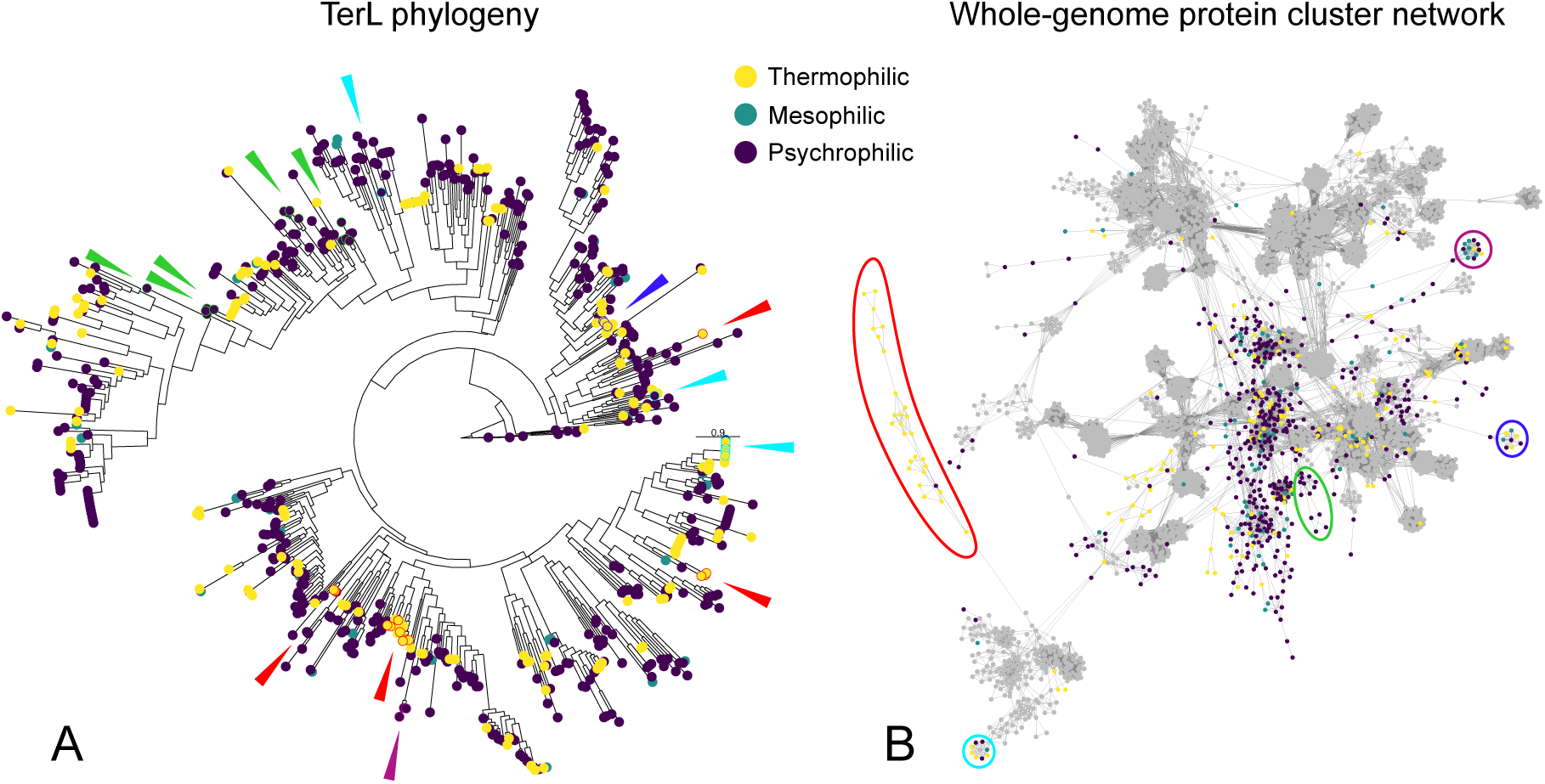
Single-gene and whole-genome comparisons of phages from enrichment cultures. **A)** Terminase large subunit (*TerL*) phylogeny (n=718). **B)** vConTACT2 whole-contig protein cluster network of reconstructed viral genomes. Grey nodes are reference sequences from Refseq release 207. The groups circled in red and magenta are largely homogenous in terms of incubation temperature, while the groups in blue, light blue, and green are highly mixed. Both figures are coloured by incubation temperature: thermophilic (≥50°C), mesophilic (<50°C and ≥ 20°C), or psychrophilic (<20°C). Groups circled in the vConTACT2 diagram in part B are labeled with arrows in the corresponding colour on the *TerL* phylogeny; not all contigs contained an annotated *TerL* gene, and thus group sizes in the protein cluster network do not correspond perfectly with the number of arrows on the phylogeny. The complete network is available in Supplementary Figure S3.

Further supporting the general lack of correlation between genome content and shared physico-chemical parameters, we found that such mixed groups were common in this dataset, with approximately 28% of viral genus-level clusters identified by vConTACT2 containing contigs assembled from enrichment cultures of different temperature groupings (psychrophilic, mesophilic, or thermophilic), and 36% containing viral contigs from different sampling locations. Despite the varied provenances and natural histories of the microorganisms in these enrichment cultures, highly related viruses were recovered across incubations. For example, viral genus-level cluster VC25 (dark blue group, Figure 3B) contained viral contigs assembled from the Elba Seep (mesophilic), Menes Caldera (mesophilic), the Black Sea (psychrophilic), Santa Monica (psychrophilic), and Guaymas Basin (thermophilic) enrichment cultures. Contigs within VC25 share multiple non-core genes, such as a *chaB* cation transport regulator (K06197) and *yfbK* Ca-activated chloride channel homolog (K07114).

These results suggest that evolutionary pressures on individual viral species/genera may have been greater before their distribution across this range of habitats, thus explaining the lack of temperature and location-related patterns at the species/genus level. Furthermore, current selection pressures may act on “accessory” genes rather than highly conserved, core genes such as *TerL* and *MCP*, resulting in some visible clustering in gene-sharing networks that is lacking in the single-gene phylogenies; these findings agree with previous reports for environmental dsDNA phages [89, 90]. However, as seen with VC25, this is not a universal rule as contigs in this cluster share many non-core genes but were assembled from enrichments of diverse provenance. When taken together with the community-level PCA results, these data may indicate that the rate of selection on viral community composition is faster than the rate of adaptation of individual viral OTUs to a specific environment.

### Viral matches to putative hosts

To link viruses with their microbial hosts in the enrichment cultures, we used CRISPR spacer matching and tRNA similarity analysis between viral genomes and potential hosts. In total, only 19 of the 231 genus-level viral clusters and an additional 16 of the unclustered viruses were matched with hosts. Notably many of these hosts belonged to low-abundance taxa within the incubation. Among the matched hosts, 13 were bacteria and 4 were archaea (Supplementary Figure S4). All represent phyla common within deep-sea alkane-rich sediments [85, 86, 91, 92]. A single CRISPR spacer match was found between a phage from the Black Sea enrichment culture and a member of the methanotrophic ANME-2c (*Ca. Methanogaster*) enriched from the Elba coastal seep, further underscoring the apparent lack of geographic constraint on the distribution of phages in such ecosystems.

Viral cluster 25, highlighted above for its presence in all enrichment cultures except for the butane-oxidizing incubation, is predicted to infect members of *Omnitrophota* (formerly OP3), a candidate phylum within the PVC superphylum, for which no phage associations have been reported to date (**Figure 4**). Characterized members of *Omnitrophota* are primarily nanobacteria from diverse environments, exhibiting genomic traits suggestive of a host-associated lifestyle [93]. Representatives of *Omnitrophota* have been previously described from seep environments and in anaerobic enrichments amended with dodecane [94] and were detected at low relative abundance (<0.5%) in several of our alkane-oxidizing enrichments. Although their functional roles in hydrocarbon-rich marine sediments remain poorly understood, the widespread recovery of VC25 across multiple sampling sites adds to our understanding of the broader ecological interactions for this lineage.

**Figure 4.**
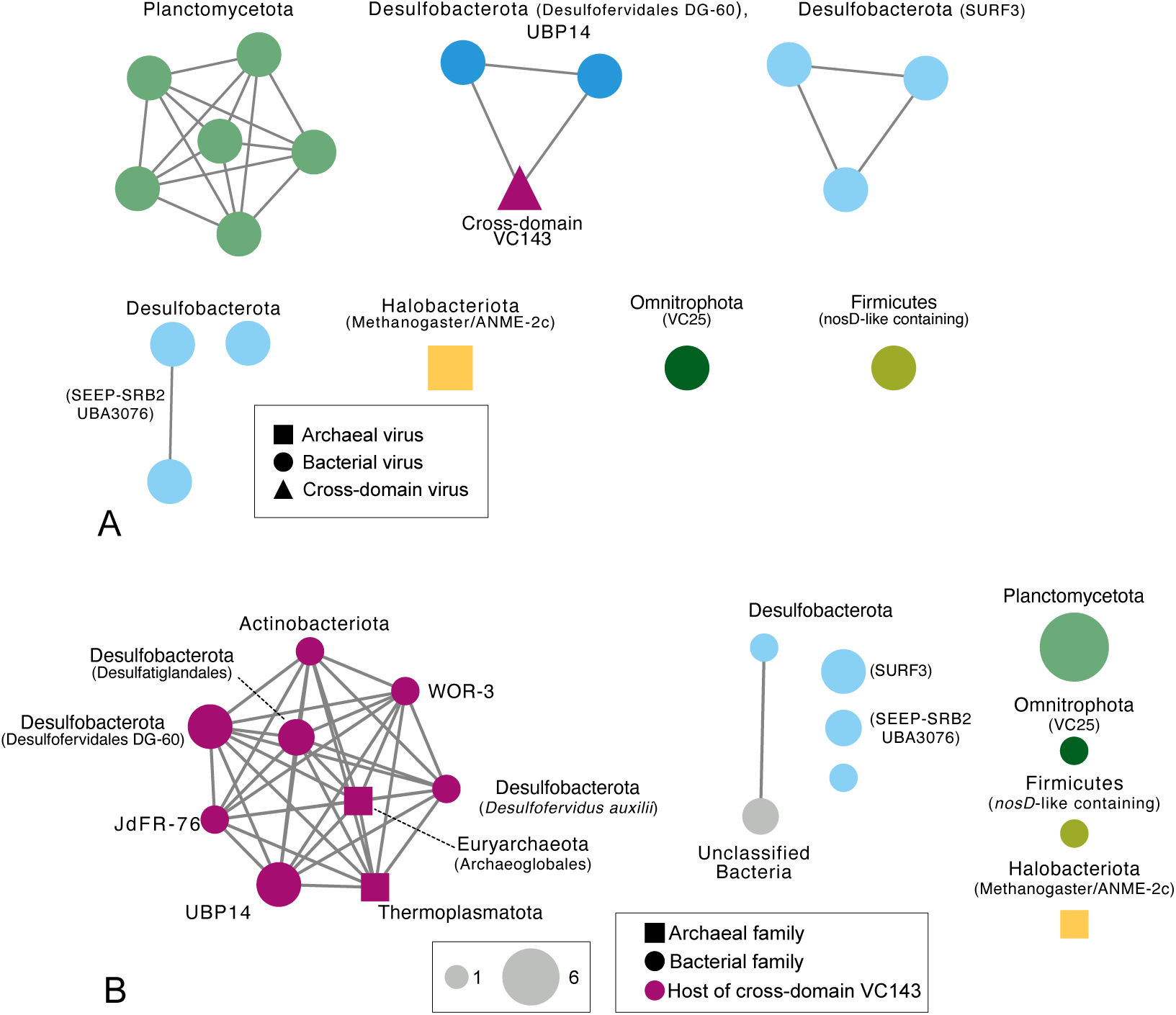
Host-virus matching. **A)** Network depicting phage genus-level clusters as nodes, and edges representing whether they share a microbial host (family-level). Nodes are labeled by microbial host phylum; with the exception of the putative cross-domain virus, any viral cluster infecting *Desulfobacterota* (putative syntrophic partners of alkane oxidizing archaea) is coloured in shades of blue. VC25 and its putative *Omnitrophota* host are coloured dark green. **B)** Network showing microbial hosts as nodes (family-level); edges represent whether the hosts are infected by the same genus-level viral cluster, and the number of edges represents the number of shared viral clusters. Nodes are labeled by host phylum. Magenta nodes represent microbial families likely infected by the broad host range viral cluster VC143, though VC143 may not be the only virus infecting the given family. The node size corresponds to the number of viral clusters matched to each family via CRISPR spacers or tRNAs. These networks have been truncated to remove most singletons and taxonomic families not addressed in the main text. Full networks are provided in Supplementary Figure S4.

While CRISPR-Cas systems are widespread across prokaryotes [95], they are nearly absent in certain lineages [96]. Furthermore, our host matching was primarily based on genomes assembled from public databases and earlier time points in these multi-year enrichment cultures. The limited matches with the most dominant microbial lineages in our alkane-oxidizing enrichment cultures may be due to high turnover rates in host CRISPR spacers [97–100] or mutation within the phage genomes in response to host immunity [101]. Indeed, comparison of community composition at the inception of certain enrichment cultures showed strong shifts in the dominant taxa by the time of this study. As an extreme example, the hydrogen-amended sulfate-reducing enrichments originally contained >95% *Desulfofervidus auxilii* (syntrophic partner of alkane-oxidizing archaea) with *Archaeoglobus sp.* as a contaminant [81]. This distribution had reversed at the time of sampling, with a low proportion of *Desulfofervidus auxilii* and a dominance of *Archaeoglobus sp*. In addition, CRISPR spacer matching as a tool for host identification is highly specific, but prone to false negatives [102]. We also note the possibility of biases in the viral community due to founder effects during the enrichment process. It is therefore unsurprising that large numbers of spacer matches were not found. However, the aforementioned match between a Black Sea enrichment culture phage and an Elba seep ANME-2c host genome suggests host-virus pairings may still be maintained despite distance and long incubation times.

### A putative broad host range virus infecting archaea and bacteria

Among the host-virus pairs identified, one viral cluster (VC143, from the 50°C Guaymas Basin methane enrichment cultures) is of particular interest due to its potential for cross-domain infections between bacteria and archaea (Figure 5). CRISPR and tRNA-identified microbial hosts of VC143 included *Desulfofervidus auxilii* (the previously characterized sulfate-reducing bacterial partner of methane-, propane-, and butane-oxidizing archaea from Guaymas Basin [15, 16, 81]), other lineages of *Desulfobacterota*, members of the archaeal *Thermoplasmatota* and *Archaeoglobus*, and several other bacterial phyla common in deep-sea thermophilic environments, including Guaymas Basin [9].

**Figure 5.**
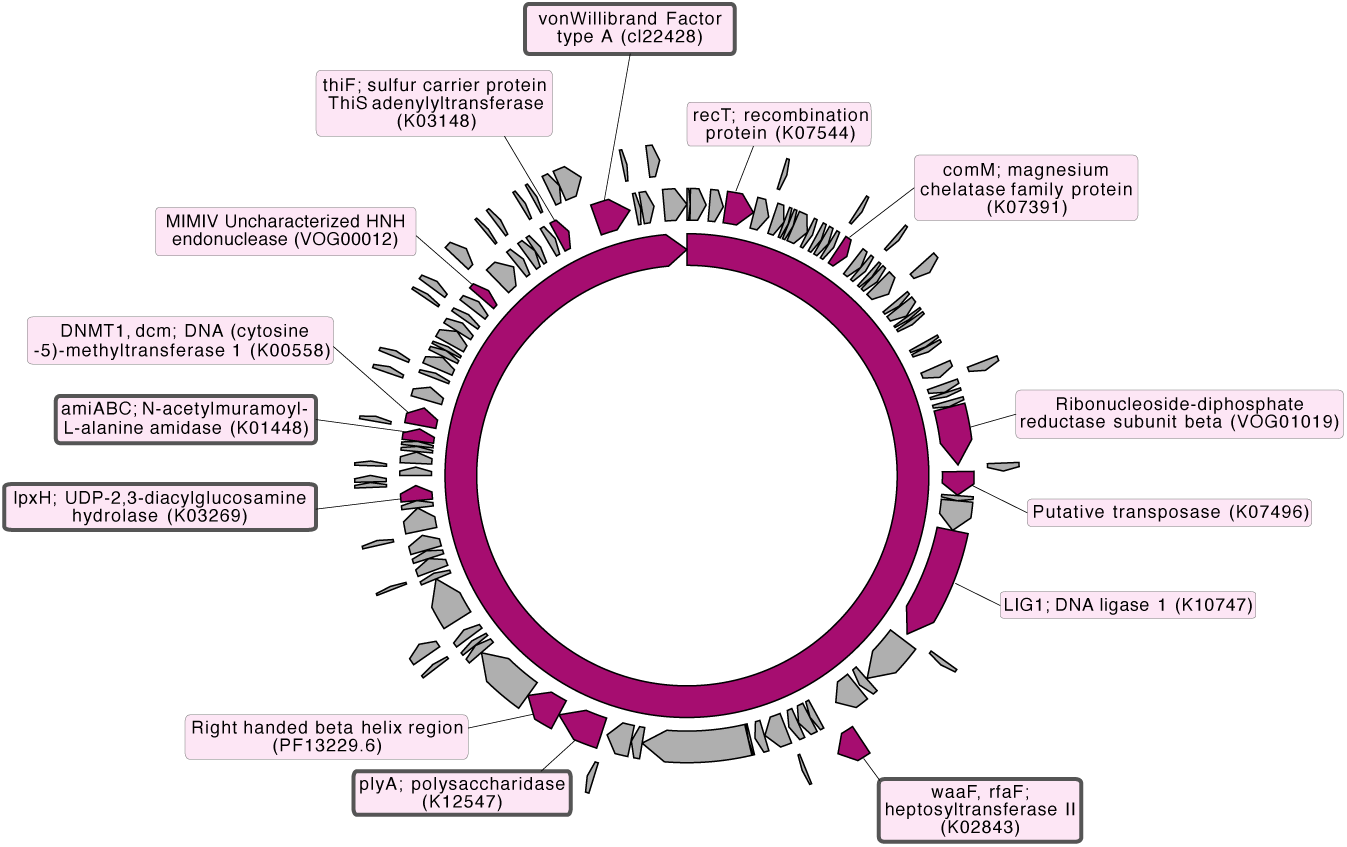
Genome map of a contig from the predicted cross-Domain infecting VC143. Annotations from either Vibrant or Conserved Domain Search are highlighted in pink, and sugar metabolism gene lab are outlined in dark grey. The grey segments are hypothetical proteins.

The two contigs in VC143, approximately 75kb in length, are longer than the average marine dsDNA phage genome size of 50kb [103, 104]. VC143 was predicted by Vibrant [51] to be a lytic virus with a circular genome. Gene markers commonly used for phylogenetic assessment such as *TerL* or *MCP* were not found in VC143, making phylogenetic placement difficult. Of the 117 predicted open reading frames in the VC143 genome, only ∼8% (14 ORFs) were annotated (identified by either Vibrant or NCBI’s Conserved Domain Search, Figure 5). These included five genes affiliated with sugar metabolism and cell wall functions (*waaF/rfaF* (K02843), a closely related vonWillibrand factor type A, *plyA* (K12547), *lpxH* (K03269), and *amiABC* (K01448)). A protein BLAST search of each of these ORFs against GTDB r214, NCBI (non-redundant protein sequences), and the cellular genomes used in this study showed similarity both to archaeal genomes and bacterial genomes across a wide number of phyla, including *Desulfobacterota*, *Methanobacteriota*, *KSB1*, and *Thermoplasmatota* (Supplementary Table S5). The diversity of sugar metabolism genes, commonly co-opted for viral adhesion to the cell surface [105], may facilitate VC143’s broad host range.

Putative cross-domain viruses have been previously reported in other methane-rich environments, including those that support biofilm formation and syntrophic interactions. An analysis of methanotrophic and methanogenic archaea found viruses predicted to infect methanogens, members of *Chloroflexi*, and members of *Deltaproteobacteria*. The cross-domain infectivity of these viruses is hypothesized to be the result of close mutualistic relationships between the host groups [27]. Additional work in hydrothermal microbial mats from Guaymas Basin used Hi-C based methods and CRISPR spacer analysis to recover viruses suspected to infect both ANME and SRB. Interestingly, the alkane-oxidizing archaeal partners of *D. auxilii* were not identified as hosts for the VC143 viruses. Targeted investigations beyond preliminary predictions from CRISPR spacer matching are necessary to robustly characterize the host range for this viral group, and it remains a possibility that the alkane-oxidizing archaea genomes used in our study may have lacked a spacer that matched the viral genomes due to CRISPR spacer turnover or changes to the phage genome in response to host defenses.

Cross-domain viral infection signatures in metagenomic datasets have been previously proposed to occur via one of four scenarios: 1) viral entry into a non-primary host or viral DNA uptake by a non-primary host, resulting spacer gain, 2) Direct DNA transfer between two organisms of different domains, including CRISPR spacer sequences, 3) Viral host switching, or 4) A true broad host range virus, capable of successful infection of both bacteria and archaea [106]. Given the breadth of prokaryotic taxa associated with the VC143 viral cluster in our dataset and the lack of spacer or tRNA matching to the archaeal partners of *D. auxilii*, scenario two is unlikely. We also predict that a complete switch in host would result in the loss of unnecessary genes, since viral genomes are highly constrained in size [107]. We caveat this proposal with the understanding that extracellular DNA is thought to be an important source of phosphorus in the deep ocean [108], and we cannot eliminate the possibility that CRISPR spacers are acquired via scavenging of extracellular viral DNA. However, we think it unlikely that DNA scavenging alone is responsible for the entry, persistence, and insertion of the spacer into the correct section of the host genome. Viruses predicted to infect both bacteria and archaea have now been found multiple times in hydrothermally influenced seep systems. Understanding whether this virus-host promiscuity extends to other sedimentary systems, including methane cold seeps, is essential to constraining their importance in horizontal gene transfer and cellular evolution in these environments.

### Auxiliary metabolic genes are not adapted to cellular metabolic lifestyles

Given the importance of methane oxidation to climate outcomes, we were particularly interested in any evidence of unique AMGs involved in alkane degradation or sulfur cycling, following the trend of reports from other sulfur-rich [109] and methane-rich [30] environments. Our dataset uniquely captures a broad range of temperatures (10-60°C) and sampling locations, providing an opportunity to examine whether the auxiliary metabolic genes (AMGs) present in virus genomes extracted from the long-term cultures were shaped by their unique environmental and incubation conditions.

Despite the extreme nature of the host metabolisms and the environments in which they are found, viral AMGs do not appear to have a great deal of specificity to hydrocarbon oxidation or sulfate reduction in these diverse enrichment cultures (Figure 6). The most abundant AMG in this dataset, annotated as DNA (cytosine-5)-methyltransferase (DNMT, K00558), was nearly ubiquitous across the assembled viromes (Figure 6) and is hypothesized to be involved in viral evasion of host defenses in various environments [110, 111].

**Figure 6.**
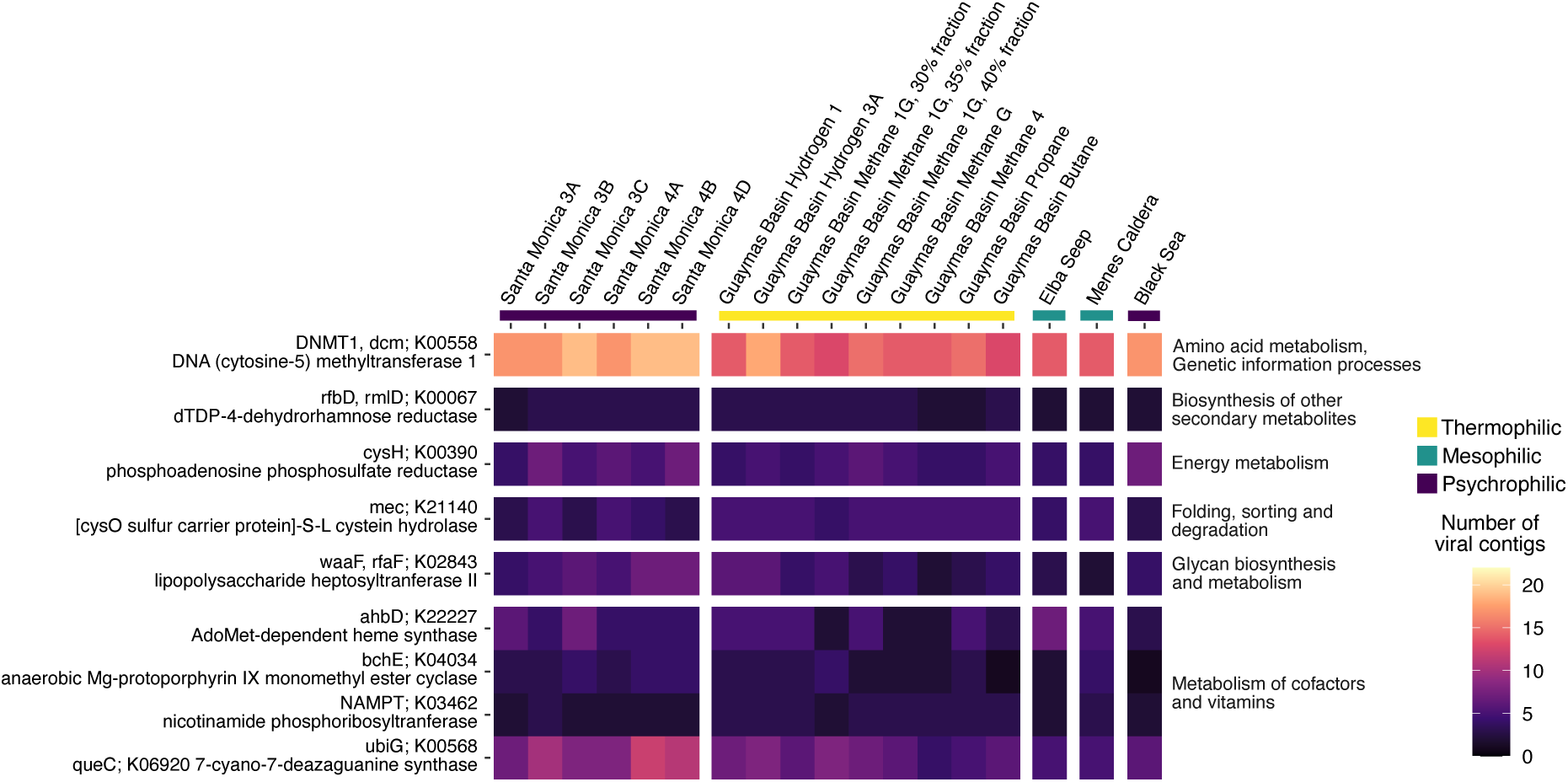
Heatmap of the number of viral contigs in each enrichment culture containing the given auxiliary metabolic gene (AMG). Contigs were included in each enrichment’s analysis only if the centered log ratio of their abundance was greater than one. AMGs were included if at least one sample had more than two contigs containing that AMG. Coloured bars under each incubation name indicate the different categories of incubation temperature. For the Guaymas Basin Methane 1G enrichment culture, we sequenced multiple fractions of viral-like particles after density gradient separation to determine if different fractions (which often enrich for different viruses) would yield variations in AMG abundances. No differences were observed. A full list of detected AMG’s the full heatmap and additional information is available in the supplement (Supplementary Figure S5).

We also found putative AMGs related to heme synthesis, which are of particular interest due to their possible role in direct interspecies electron transfer (DIET) [112–114]. DIET is hypothesized to underlie the syntrophic archaeal/bacterial partnership [15, 115–117], in which electrons from the alkane-oxidizing archaea are transferred directly to the bacterial partner for sulfate reduction. Heme-related AMGs in this study include heme synthase (*ahbD*; K22227) and anaerobic Mg-protoporphyrin IX monomethyl ester cyclase (*bchE*; K04034). *ahbD* and its homologs are known for their roles in the biosynthesis of heme via siroheme-dependent pathways in methanogenic and methanotrophic archaea and sulfate-reducing bacteria [118–121], and *bchE* is involved in chlorophyll synthesis in anaerobic phototrophs, though protoporphyrin IX is also an intermediate in the protoporphyrin-dependent heme biosynthesis pathway [122].

Viral replication may be enhanced through manipulation of heme-dependent respiratory processes found in both the ANME and SRB; AMGs like *ahbD* [123], *bchE* [124], and other radical SAM proteins may influence host energy availability via manipulation of cellular redox states. While hosts of the viruses carrying these genes have not been identified and biochemical and phylogenetic characterization is necessary, the high demand for both iron and sulfur by alkane-oxidizing archaea and their partner sulfate-reducing bacteria [125] and their reliance on heme for energy generation might offer clues as to which organisms are most susceptible to viral manipulation with *ahbD* and *bchE*.

The majority of the AMGs found, including *ahbD* and *bchE*, play general roles in the biosynthesis of amino acids and cofactors, are broadly important for the maintenance of cellular metabolism, and have been found in a variety of systems, including methane cold seep sediments [28, 29]; cow rumen [126]; compost systems [127]; Baltic Sea sediments and water column [123]; the millipede gut [124]; and a variety of marine, freshwater, terrestrial, and engineered systems [109]. AMGs directly related to hydrocarbon oxidation or sulfate reduction were not detected, although methane oxidation-related AMGs (e.g. *pmoC*) have been previously found in freshwater systems [30] and sulfate metabolism AMGs (e.g. *dsrA*) are abundant in a variety of aqueous environments [109, 128]. Perhaps the selection pressure exerted by the need to find a host in a compact, heterogeneous sediment matrix is greater than the competitive advantage available from specializing to a particular host metabolic lifestyle.

### *nosD*-like proteins likely involved in adhesion

We detected several viral genes annotated by Vibrant as nitrous oxide reductase subunit D (*nosD*), as well as additional genes annotated as sugar metabolism genes that, according to NCBI’s Conserved Domain Search, shared similarities to *nosD*. As nitrous oxide reductase plays an important role in bacterial denitrification and nitrogen cycling more broadly, we investigated the distribution and potential function of the *nosD* homolog among the viruses in the alkane enrichment cultures. A custom *nosD* HMM search, in conjunction with Vibrant annotations, uncovered *nosD*-like genes in four genus-level vMAG clusters (VC18, VC77, VC239, VC276) and two additional unclustered contigs. vMAGs containing *nosD*-like genes were assembled across multiple enrichment cultures: Santa Monica Basin (10°C), Guaymas Basin (methane, propane, and butane incubations, all 50°C), and the Black Sea (12°C). An additional search of viral contigs from Costa Rica methane seep sediments dominated by ANME-2 subgroups and their SRB partners also uncovered several *nosD*-like genes [72] (Figure 7A).

**Figure 7.**
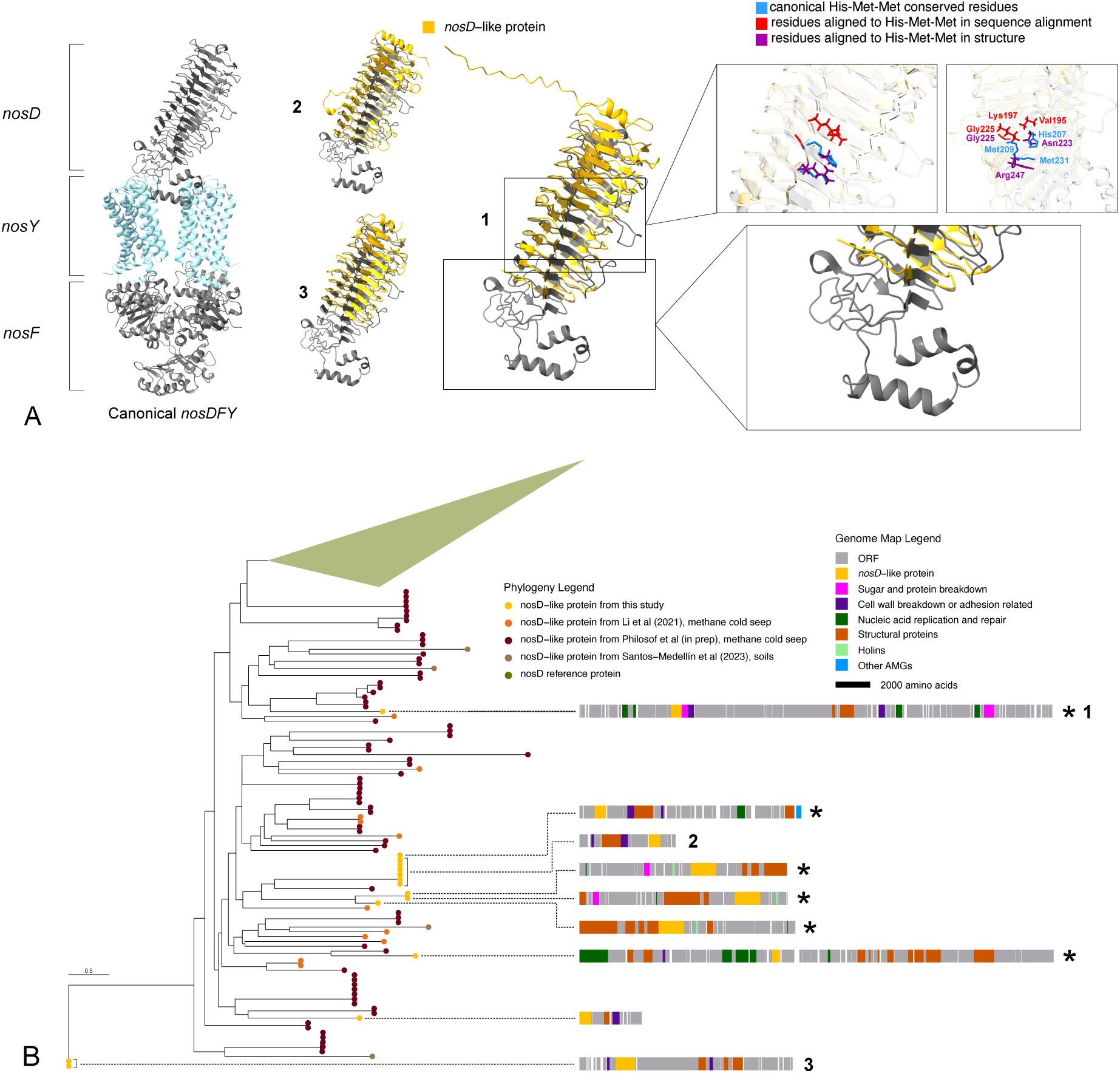
Structural and phylogenetic analysis of nosD-like protein. **A)** Examples of modeled structures of viral *nosD*-like proteins from this study, shown in yellow and overlaid on the canonical *nosD* (dark grey). On the left is the canonical *nosDFY* structure. Insets on the right highlight the two major structural differences between these nosD-like proteins and the canonical protein. The upper inset panels show the loss of the His-Met-Met motif, with the red residues corresponding to the motif in the sequence alignment, and the purple residues corresponding to the motif in the structural alignment. Gly225 corresponds to an active site residue in both the sequence and structure alignments. The lower panel shows the loss of the structures necessary to bind to the *nosY* subunit, suggesting an alternative role for the *nosD*-like proteins. Structures of the full complement of *nosD*-like proteins from this study are provided in Supplementary Figure S7, and those referenced in Figure 6B are available from the CaltechDATA (See Data Availability). **B)** Phylogeny of *nosD*-like proteins from this study (found using hmmsearch or from Vibrant annotations) and mined from other methane cold seep datasets [28, 72] and terrestrial soils [74]. The green clade consists of known canonical *nosD* proteins. *nosD*-like proteins from this study are connected by dotted lines to a gene map of all open reading frames found in the assembled contig (in some cases, this does not represent the full length of the assembled contig as ORFs were not necessarily found across the entire length); categorization of proteins besides the *nosD*-like sequences are based on Vibrant annotations. Approximate length scale of genomes is shown under the Genome Map Legend. Asterisks indicate contigs where the Vibrant-called ORF containing the *nosD*-like protein also contains additional adhesion or sugar breakdown related domains. The numbered genomes (1, 2, and 3) correspond to the structures labeled in part A.

Canonical NosD protrudes into the periplasm and is anchored to the NosY inner membrane subunit, where it shuttles a copper ion to the active site of NosZ [70]. A highly conserved histidine-methionine-methionine motif serves as a transient copper binding site that may also facilitate sulfur shuttling [70, 129]. Several viral-encoded *nosD*-like sequences (filtered at >200 amino acids) spanned the conserved active site of the canonical NosD domain, but appeared to have lost the His-Met-Met motif and lacked the structural components required for NosY interaction (Figure 7A, Supplementary Figure S6).

The absence of characteristics central to the canonical function of NosD suggests an alternative function for these NosD-like proteins. The NosD domain has known similarities to the CASH domain, which is a widespread carbohydrate and sugar binding domain with homologs to archaeal S-layer proteins [105]. The three-sided right-handed beta-helix fold in NosD, involved in glycan interactions like depolymerization, is also thought to potentially function in a glycan binding role [130]. In several of the viral contigs in which NosD-like proteins are found, these proteins share an ORF with attachment and sugar depolymerization domains that are typically extracellular such as pectate lyase and concanavalin A type lectin/glucanase (Figure 7B). Other ORFs on these viral contigs encode additional cell wall and sugar related proteins such as glycosyltransferases involved in cell wall biosynthesis and mannosyltransferases (annotated by Vibrant and confirmed using NCBI Conserved Domain Search). Prior work has hypothesized that AMGs related to O-antigen synthesis in viruses of methanotrophic and methanogenic archaea are important for enhancing cell-cell adhesion, potentially resulting in both improved host survival rates via aggregation and higher probability that the phage will encounter another host [27]. Such genes, including *galE* (K01784) and *rfbD* (K00067), were also present in viruses from our enrichments (Figure 6 and Supplementary Figure S5). We hypothesize that viral *nosD*-like genes have a similarly important role related to cell-surface binding and sugar metabolism.

One notable contig (labeled “1” in Figure 7) contained a NosD-like protein that shares its ORF with a dockerin domain. Other portions of the contig encoded an archaeal S-layer protein and additional dockerin domains. Dockerin domains have been previously described in ANME-1 and ANME-2 genomes [131, 132] and are predicted to be involved in cell-cell adhesion and attachment of proteins such as transporters to the outside of the cell. These genes were most closely related to genes from ANME-2 genomes (BLASTp, NCBI non-redundant protein sequences database), suggesting this may be a virus capable of infecting these methanotrophic archaea. ANME-2 archaea frequently occur in tightly packed syntrophic consortia with sulfate-reducing bacteria and the presence of predicted cell-surface genes in this virus near the *nosD*-like gene is consistent with the predicted role in adhesion. Another notable *nosD*-like-containing viral cluster putatively infects members of the bacterial class *Desulfotomaculia* (phylum *Firmicutes*, Figure 4). These observations further suggest that viruses encoding *nosD*-like domains are associated with diverse microbial hosts.

The *nosD*-like genes found in these enrichments are variably located in the assembled viral genomes, with some located within areas of high AMG concentration and many located near viral structural and assembly genes such as terminases, prohead proteases, and tail tube proteins (Figure 7A); this suggests that NosD-like proteins take on diverse functions that likely vary among phages. We propose three possible roles for virus-encoded NosD-like proteins: 1) *nosD*-like proteins are involved in viral contact and adhesion to the host at the start of infection, 2) *nosD*-like proteins are involved in lysis of the host cell at the end of infection via adhesion to and breakdown of the peptidoglycan layer and/or extracellular matrix, and 3) *nosD*-like proteins are active during infection and influence adhesion between the host and other cells (Figure 8). The first and second hypotheses are supported by the presence of homologous right-handed beta-helix domains in both *nosD* and phage tailspike proteins, the latter of which are generally polysaccharide lyases and glycosyl hydrolases [130].

**Figure 8.**
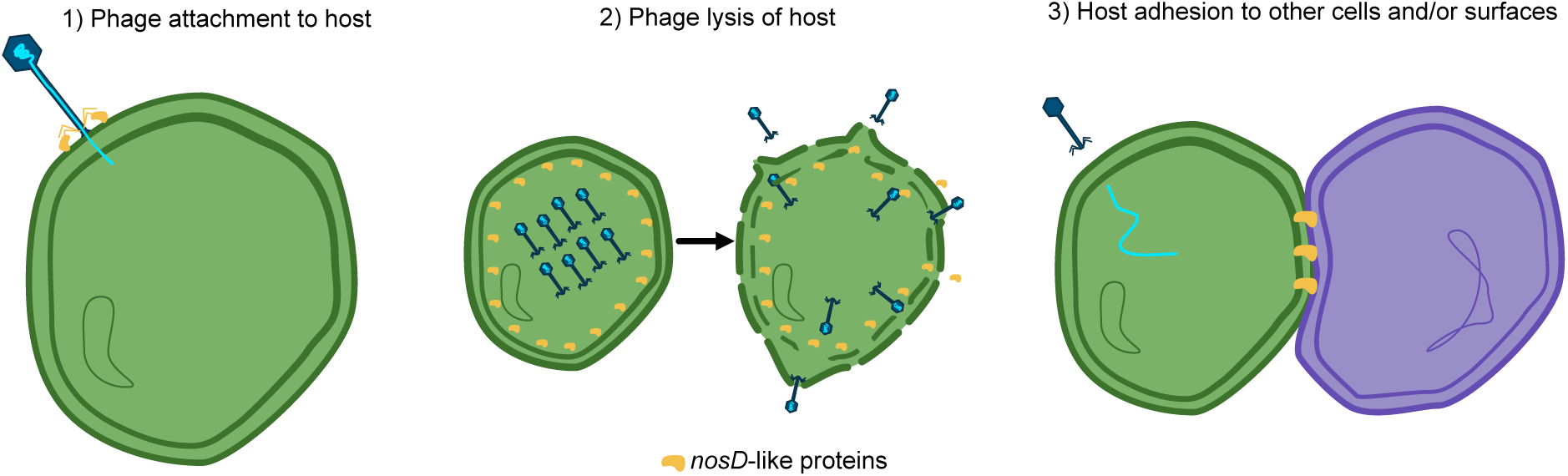
Schematic of the three hypotheses for the role of these *nosD*-like proteins. In hypothesis 1, *nosD*-like proteins are involved in the attachment of the phage to the host at the very beginning of infection. In 2, *nosD*-like proteins are expressed near the end of infection to assist with lysis of the host via breakdown of the cell wall. In 3, *nosD*-like proteins play a greater role in cell physiology, influencing host tendency and ability to attach to other cells and surfaces.

Additional support for all three hypotheses comes from the absence of evidence for these longer *nosD*-like proteins in water column datasets. Our search of viral genomes from the Tara Oceans Project [71] using our *nosD* HMM identified several shorter hits, but none exceeded ∼200 amino acids or reliably spanned the conserved copper-shuttling motif in a multiple sequence alignment against canonical *nosD* proteins. We also examined both sediment and water column phages from the meromictic lake Lac Pavin [73]; and while all hits were shorter than 200 amino acids, they were exclusively found within phages from the sediment. Searches of additional datasets from a previous methane seep study [28] and terrestrial soils [74] also yielded putative *nosD*-like proteins (Figure 7B). As microbes in sediment and soil ecosystems have more opportunities for surface attachment than those in the water column, viruses in sediments may benefit from additional sugar breakdown and adhesion proteins for host infection within complex extracellular matrices (matching previous work in biofilms [133]), or they may carry *nosD*-like proteins to influence their hosts’ adhesion to other cells and substrates.

## Conclusions and Future Directions

This study highlights the advantages of using sediment-free enrichment incubations to study the viruses in a variety of slow-growing, hydrocarbon-oxidizing systems. Removing the sediment matrix and selecting for microorganisms of interest based on temperature and substrate greatly reduced the complexity inherent in deep-sea sediment communities, allowing for targeted investigations. We found that despite variation in temperature regimes, sampling locations, and energy sources, viral genomes showed no consistent patterns in phylogeny or gene content.

While the effects of incubation temperature and original sampling location cannot be deconvolved, these parameters may instead have a greater influence in structuring viral community composition.

The discovery of a putative cross-domain virus in this system adds to existing knowledge of cross-domain phages in sediment and biofilm-dominated environments. This raises the question of whether cross-domain phages are more common in low-energy sediments than they are in the water column. In the latter, the odds of encountering a host are likely higher and high-energy yield host metabolisms enable faster reproduction, thus potentially generating greater host specificity. Isolation of such phages coupled to long-term evolutionary experiments and targeted analysis of existing phage datasets are necessary for further testing of these hypotheses.

Since our current understanding of marine phages is strongly biased towards the upper water column, working in sediment communities is crucial to understanding the role phages play in these important carbon sinks. The presence of unique *nosD*-like proteins in these sediment-derived enrichments, sediment incubations from methane cold seeps, and terrestrial soils, but their conspicuous absence in water column datasets, supports evidence for novel adhesion roles in communities dominated by aggregation and attachment-based lifestyles. Further investigation of *nosD*-like proteins would be greatly aided by their expression in genetically tractable model systems followed by biochemical characterization. Such investigations may prove interesting to the medical field, where an estimated 80% of human bacterial infections are thought to be biofilm-dominated [134, 135]; searching for *nosD*-like proteins in viral communities of such infections could shed light on the dynamics of biofilm construction and chronic infection.

Taken together, the presence of viruses infecting both archaea and bacteria, the lack of AMGs related to core host metabolic processes, and the presence of *nosD*-like proteins suggest that viruses may be more specialized to the complexity of the sediment matrix and the resulting presence of surface-attached hosts, rather than to other physicochemical parameters like temperature or specific host metabolic processes.

The presence of viruses in these sediment-free enrichments serves as a reminder that our standard techniques for understanding laboratory or environmental communities may yield an incomplete picture. As the field of microbial ecology develops more powerful techniques to enrich and isolate organisms of interest, we should be careful not to overlook the possible contributions of viruses to enrichment composition and stability.

## Supporting information

Narayanan_etal_2025_supp

## Acknowledgements

The authors would like to acknowledge the following individuals and groups who helped make this work possible. From the University of Southern California, Dr. Zarko Manojlovic and Ashley Jauregui for performing the metaviromic sequencing; Dr. Daan R. Speth and Dr. Rafael Laso-Pérez for allowing us to use their unpublished genomes in our hunt for host-virus pairs; Dr. Igor Antoshechkin at the Millard and Muriel Jacobs Genetics and Genomics Laboratory at Caltech for library preparation and sequencing of the cellular metagenomes; Dr. Hang Yu for initial incubation development from Santa Monica and long-term maintenance of other laboratory enrichments used in this study; Anja Seebeck and Susanne Menger at the University of Bremen for help with cultivation and incubation maintenance; Dr. Daniel Utter for bioinformatics advice, Dr. Justus W. Fink and Dr. Jeremy E. Schreier for manuscript feedback; and the shipboard scientists and crews of the R/V Poseidon exp. 317/3, R/V Atlantis AT15-45 and AT37-06, RV L’Atalante NAUTINIL Expedition, and the MBARI RV Western Flyer 2013 Southern California Expedition; Hydra Station in Fetovaia, Italy for seep sample collection and curation.

## Data availability

Raw fastqs from the metagenome, metavirome, and 16S rRNA sequencing are under NCBI PRJNA1226641. All MAGs generated in this study for which a virus CRISPR or tRNA match was found, all 989 concatenated viral contigs and associated metadata, viral protein annotations, *nosD*-like sequences used in Figure 7, pdb files of all *nosD*-like sequences from Figure 7, and the *nosD* HMM file are available at https://doi.org/10.22002/7sp3k-ewh62.

