## Supplementary material for "Viral communities from long-term anaerobic alkane-oxidizing enrichments may promote cell surface adhesion": Narayanan_etal_2025_supp

### Supplementary information

| Sample | Sampling Lat, Long | Incubation Temp. (˚C) | Energy substrate | Year of collection | Year of incubation start | Supplementary references for media recipe |
|---|---|---|---|---|---|---|
| Santa Monica 3A | 33.78888889, -118.6680556 | 10 | 0.2MPa Methane | 2013 | 2013 | [1] |
| Santa Monica 3B | 33.78888889, -118.6680556 | 10 | 0.2MPa Methane | 2013 | 2013 | [1] |
| Santa Monica 3C | 33.78888889, -118.6680556 | 10 | 0.2MPa Methane | 2013 | 2013 | [1] |
| Santa Monica 4A | 33.78888889, -118.6680556 | 10 | 0.2MPa Methane | 2013 | 2013 | [1] |
| Santa Monica 4B | 33.78888889, -118.6680556 | 10 | 0.2MPa Methane | 2013 | 2013 | [1] |
| Santa Monica 4D | 33.78888889, -118.6680556 | 10 | 0.2MPa Methane | 2013 | 2013 | [1] |
| Guaymas Basin Hydrogen 1A | 27.01213889, -111.4152222 | 60 | 0.2MPa 80:20 Hydrogen:$CO_2$ | 2009 | 2012 | [2], Extended Methods from this study |
| Guaymas Basin Hydrogen 3 | 27.01213889, -111.4152222 | 60 | 0.2MPa 80:20 Hydrogen:$CO_2$ | 2009 | 2012 | [2], Extended Methods from this study |
| Guaymas Basin Methane 1G | 27.01666667, -111.6833333 | 50 | 0.2MPa Methane | 2009 | 2010 | [3], Extended Methods from this study |
| Guaymas Basin Methane G | 27.01666667, -111.6833333 | 50 | 0.2MPa Methane | 2009 | 2010 | [3], Extended Methods from this study |
| Guaymas Basin Methane 4 | 27.01666667, -111.6833333 | 50 | 0.2MPa Methane | 2009 | 2010 | [3], Extended Methods from this study |
| Guaymas Basin Propane | 27.01213889, -111.4152222 | 50 | 0.15MPa Propane | 2009 | 2010 | [4] |
| Guaymas Basin Butane | 27.01213889, -111.4152222 | 50 | 0.15MPa Butane | 2009 | 2010 | [4] |
| Elba Seep | 42.73507778, 10.2 | 20 | 0.2MPa Methane | 2010 | 2010 | [5] |
| Menes Caldera 2 | 32.30091667, 28.43519444 | Room temperature | 0.2MPa Methane | 2003 | 2003 | [6] |
| Menes Caldera 2A | 32.30091667, 28.43519444 | Room temperature | 0.2MPa Methane | 2003 | 2003 | [6] |
| Black Sea | 44.77777778, 31.97222222 | 12 | 0.2MPa Methane | 2000 | 2004 | [6] |

**Table S1.** Incubation bottle metadata. All bottles include sulfate as an electron acceptor. A full description of how the enrichment cultures (excepting Santa Monica) were set up is contained in Supplementary Reference #[7]. Santa Monica enrichments are described in Supplementary Reference #[1].

| Sample | Organism (Family level) | Relative Abundance (Family level) | Number of ASVs | ASVs comprising >1% of reads in the sample (% total reads in sample) |
|---|---|---|---|---|
| Santa Monica 3A | Fermentibacteraceae | 17.4% | 13 | ASV_16 (13.6%), ASV_15 (1.4%) |
| | ANME-2a-2b | 16.8% | 17 | ASV_10 (14.7%) |
| | Fusibacteraceae | 16.5% | 7 | ASV_40 (14.1%) |
| | ANME-2c | 9.9% | 2 | ASV_12 (8.1%), ASV_81 (1.8%) |
| | Dissulfuribacteraceae | 6.0% | 8 | ASV_25 (4.1%) |
| | Lentimicrobiaceae | 5.3% | 13 | ASV_7 (4.8%) |
| | AKAU3564 sediment group | 5.0% | 20 | ASV_193 (1.6%), ASV_224 (1.2%) |
| | Marinilabiliaceae | 4.7% | 6 | ASV_50 (3.4%) |
| Santa Monica 3B | ANME-2a-2b | 27.1% | 9 | ASV_10 (19.8%), ASV_6 (6.1%) |
| | OM190 | 18.9% | 5 | ASV_32 (11.8%), ASV_22 (2.5%), ASV_124 (2.1%), ASV_128 (2.0%) |
| | Marinilabiliaceae | 11.3% | 10 | ASV_43 (7.2%), ASV_50 (3.4%) |
| | JS1 | 9.2% | 7 | ASV_14 (5.4%), ASV_58 (2.8%) |
| | Fermentibacteraceae | 8.5% | 7 | ASV_15 (5.8%), ASV_31 (2.3%) |
| | Desulfosarcinaceae | 6.6% | 18 | ASV_45 (3.3%), ASV_147 (1.2%) |
| Santa Monica 3C | Lentimicrobiaceae | 45.0% | 30 | ASV_7 (43.9%) |
| | ANME-2a-2b | 26.3% | 6 | ASV_6 (25.7%) |
| | Fermentibacteraceae | 10.6% | 11 | ASV_15 (9.7%) |
| | Anaerolineaceae | 3.1% | 12 | ASV_86 (2.9%) |
| | Desulfosarcinaceae | 3.1% | 18 | ASV_45 (1.9%) |
| Santa Monica 4A | ANME-2c | 31.0% | 7 | ASV_12 (16.7%), ASV_74 (6.7%), ASV_77 (6.3%) |
| | Fermentibacteraceae | 9.8% | 7 | ASV_31 (9.6%) |
| | OM190 | 9.7% | 5 | ASV_22 (8.9%) |
| | Unclassified Methanosarciniales | 8.4% | 8 | ASV_57 (8.2%) |
| | Marine Benthic Group D and DHVEG-1 | 5.7% | 11 | ASV_49 (4.0%) |
| | Lokiarchaeia | 3.7% | 11 | ASV_60 (3.5%) |
| | Desulfosarcinaceae | 3.6% | 10 | ASV_114 (2.2%) |
| | Dissulfuribacteraceae | 3.6% | 4 | ASV_25 (2.3%) |
| | Unclassified Phycisphaerae | 3.2% | 6 | ASV_110 (2.8%) |
| Santa Monica 4B | Dissulfuribacteraceae | 19.0% | 6 | ASV_25 (8.7%), ASV_27 (3.6%), ASV_79 (3.4%), ASV_82 (3.2%) |
| | ANME-2c | 18.0% | 5 | ASV_12 (15.5%), ASV_81 (2.1%) |
| | Fermentibacteraceae | 13.0% | 17 | ASV_16 (11.8%) |
| | ANME-2a-2b | 8.5% | 21 | ASV_10 (3.7%), ASV_6 (3.4%) |
| | OM190 | 6.5% | 4 | ASV_22 (5.4%) |
| | Lentimicrobiaceae | 5.5% | 19 | ASV_7 (1.2%) |

| | | | | |
|---|---|---|---|---|
| | Marine Benthic Group D and DHVEG-1 | 3.8% | 22 | ASV_49 (1.3%) |
| | JS1 | 3.6% | 5 | ASV_14 (2.9%) |
| Santa Monica 4D | Dissulfuribacteraceae | 18.4% | 8 | ASV_25 (5.8%), ASV_27 (5.7%), ASV_79 (3.5%), ASV_82 (3.3%) |
| | ANME-2c | 11.6% | 5 | ASV_12 (9.0%), ASV_81 (1.7%) |
| | ANME-2a-2b | 10.4% | 17 | ASV_10 (5.0%), ASV_6 (3.3%) |
| | Marine Benthic Group D and DHVEG-1 | 9.3% | 21 | ASV_49 (2.8%), ASV_28 (1.2%) |
| | Fermentibacteraceae | 8.2% | 11 | ASV_16 (7.5%) |
| | Lentimicrobiaceae | 8.0% | 21 | ASV_7 (2.8%), ASV_164 (1.0%) |
| | OM190 | 5.7% | 4 | ASV_22 (4.8%) |
| | JS1 | 4.1% | 5 | ASV_14 (3.3%) |
| | AKAU3564 sediment group | 3.3% | 17 | None (all ASVs <1%) |
| Menes Caldera 2 | Lentimicrobiaceae | 28.7% | 7 | ASV_17 (27.6%) |
| | Marine Benthic Group D and DHVEG-1 | 25.6% | 27 | ASV_28 (9.2%), ASV_69 (6.4%), ASV_80 (4.0%), ASV_67 (2.0%), ASV_89 (1.6%) |
| | Fermentibacteraceae | 7.2% | 4 | ASV_39 (7.1%) |
| | Anaerolineaceae | 5.0% | 22 | ASV_64 (2.9%) |
| | Dissulfuribacteraceae | 4.8% | 7 | ASV_87 (4.2%) |
| | JS1 | 4.4% | 5 | ASV_14 (3.7%) |
| | Calditrichaceae | 3.5% | 6 | ASV_41 (1.5%), ASV_150 (1.2%) |
| Menes Caldera 2A | ANME-2a-2b | 25.7% | 9 | ASV_36 (15.6%), ASV_10 (4.7%), ASV_100 (3.2%), ASV_184 (1.3%) |
| | Marine Benthic Group D and DHVEG-1 | 21.2% | 26 | ASV_28 (7.2%), ASV_67 (3.1%), ASV_89 (2.3%), ASV_80 (2.2%), ASV_131 (1.3%), ASV_49 (1.1%) |
| | Unclassified Plactomycetota | 9.3% | 3 | ASV_68 (8.6%) |
| | Lentimicrobiaceae | 9.0% | 3 | ASV_85 (5.2%), ASV_17 (3.8%) |
| | Fermentibacteraceae | 7.4% | 4 | ASV_39 (7.4%) |
| | Anaerolineaceae | 7.3% | 18 | ASV_64 (6.4%) |
| | Calditrichaceae | 5.5% | 5 | ASV_41 (5.1%) |
| | JS1 | 4.4% | 4 | ASV_14 (4.2%) |
| Black Sea | JS1 | 23.0% | 14 | ASV_24 (17.1%), ASV_14 (3.7%) |
| | Desulfosarcinaceae | 18.7% | 21 | ASV_23 (18.4%) |
| | ANME-1b | 13.4% | 2 | ASV_35 (13.2%) |
| | Anaerolineaceae | 7.6% | 29 | ASV_54 (6.2%) |
| | Pirellulaceae | 7.0% | 24 | ASV_91 (4.6%) |
| | Fermentibacteraceae | 5.6% | 23 | ASV_16 (1.8%), ASV_202 (1.1%) |
| | ANME-1 | 4.4% | 12 | ASV_3 (4.2%) |
| | ANME-2a-2b | 3.6% | 15 | ASV_133 (2.3%) |

| | | | | |
|---|---|---|---|---|
| Elba Seep | ANME-1a | 44.7% | 1 | ASV_5 (44.7%) |
| | Dissulfuribacteraceae | 10.1% | 11 | ASV_33 (6.3%), ASV_27 (3.3%) |
| | AKAU3564 sediment group | 7.1% | 11 | ASV_38 (5.9%) |
| | Desulfosarcinaceae | 5.2% | 11 | ASV_53 (4.2%) |
| | Unclassified Bacteria | 5.1% | 7 | ASV_44 (5.0%) |
| | ANME-2c | 4.4% | 1 | ASV_51 (4.4%) |
| | ANME-2a-2b | 3.6% | 2 | ASV_66 (3.6%) |
| Guaymas Basin Methane 1G | Bathyarchaeia | 34.2% | 30 | ASV_8 (17.1%), ASV_9 (14.8%) |
| | ANME-1 | 18.8% | 12 | ASV_3 (15.8%), ASV_47 (2.8%) |
| | Aerophobales | 16.5% | 7 | ASV_13 (16.3%) |
| | Desulfofervidaceae | 7.9% | 11 | ASV_30 (4.1%), ASV_2 (3.1%) |
| | Thermodesulfobacteriaceae | 7.5% | 2 | ASV_19 (7.5%) |
| | GN01 | 3.0% | 1 | ASV_56 (3.0%) |
| Guaymas Basin Methane G | Bathyarchaeia | 35.9% | 24 | ASV_8 (17.8%), ASV_9 (10.8%), ASV_55 (5.0%) |
| | Desulfofervidaceae | 13.2% | 12 | ASV_2 (7.2%), ASV_30 (5.0%) |
| | Aerophobales | 12.2% | 8 | ASV_26 (10.1%), ASV_13 (1.7%) |
| | ANME-1 | 7.4% | 4 | ASV_3 (4.3%), ASV_47 (3.1%) |
| | Thermodesulfobacteriaceae | 6.1% | 2 | ASV_19 (5.8%) |
| | Aminicenantales | 5.0% | 4 | ASV_72 (2.8%), ASV_11 (2.1%) |
| | Caldatribacteriaceae | 3.6% | 1 | ASV_76 (3.6%) |
| Guaymas Basin Methane 4 | ANME-1 | 53.8% | 16 | ASV_3 (52.1%) |
| | Desulfofervidaceae | 29.0% | 13 | ASV_2 (28.2%) |
| | Bathyarchaeia | 10.3% | 24 | ASV_9 (4.7%), ASV_8 (4.5%) |
| Guaymas Basin Hydrogen 1 | Archaeoglobaceae | 93.3% | 3 | ASV_1 (93.0%) |
| | Desulfofervidaceae | 4.5% | 1 | ASV_2 (4.5%) |
| Guaymas Basin Hydrogen 3A | Archaeoglobaceae | 74.9% | 2 | ASV_1 (74.8%) |
| | Desulfofervidaceae | 22.7% | 5 | ASV_2 (22.5%) |
| Guaymas Basin Propane | Desulfofervidaceae | 25.3% | 22 | ASV_2 (23.3%) |
| | Syntrophoarchaeaceae | 13.6% | 6 | ASV_20 (13.3%) |
| | JS1 | 8.7% | 4 | ASV_21 (8.6%) |
| | Marine Benthic Group D and DHVEG-1 | 4.1% | 5 | ASV_102 (2.2%), ASV_144 (1.2%) |
| Guaymas Basin Butane | Aminicenantales | 23.4% | 4 | ASV_11 (23.4%) |
| | Desulfofervidaceae | 9.9% | 9 | ASV_2 (9.6%) |
| | Syntrophoarchaeaceae | 3.7% | 2 | ASV_107 (2.2%), ASV_20 (1.5%) |
| | JS1 | 3.4% | 3 | ASV_21 (3.4%) |

**Table S2.** ASVs comprising at least 1% of total reads in the given sample, with their family-level classification. Only families >3% relative abundance are shown. The majority of families are comprised of no more than two ASVs.

| | Richness | | Simpson's Evenness | | EQ | | Evar | |
|---|---|---|---|---|---|---|---|---|
| **Sample** | **Viral** | **Cellular** | **Viral** | **Cellular** | **Viral** | **Cellular** | **Viral** | **Cellular** |
| Santa Monica 3A | 313 | 259 | 0.00326246 | 0.05160458 | 0.06033558 | 0.10089224 | 0.0733037 | 0.18937416 |
| Santa Monica 3B | 395 | 230 | 0.00711382 | 0.05803197 | 0.05029099 | 0.0912637 | 0.04945798 | 0.15952748 |
| Santa Monica 3C | 384 | 209 | 0.01598118 | 0.01766819 | 0.04877084 | 0.09484208 | 0.04572982 | 0.18153026 |
| Santa Monica 4A | 412 | 260 | 0.0155338 | 0.0565383 | 0.0547177 | 0.0957181 | 0.05905509 | 0.18104189 |
| Santa Monica 4B | 422 | 363 | 0.01737859 | 0.04896974 | 0.0504078 | 0.1093796 | 0.04927744 | 0.23193287 |
| Santa Monica 4D | 417 | 342 | 0.02387932 | 0.08732644 | 0.05097117 | 0.11283316 | 0.05002626 | 0.23445262 |
| Guaymas Basin Hydrogen 1 | 333 | 40 | 0.03758998 | 0.0287957 | 0.06136846 | 0.07591399 | 0.07500451 | 0.11558463 |
| Guaymas Basin Hydrogen 3A | 319 | 66 | 0.03382399 | 0.02478285 | 0.06205792 | 0.08885116 | 0.07720127 | 0.20135376 |
| Guaymas Basin Methane 1G, 30% optiprep | 331 | N/A | 0.00413344 | N/A | 0.05902468 | N/A | 0.06943501 | N/A |
| Guaymas Basin Methane 1G, 35% optiprep | 325 | N/A | 0.00424544 | N/A | 0.05218896 | N/A | 0.0567324 | N/A |
| Guaymas Basin Methane 1G, 40% optiprep | 320 | N/A | 0.00935381 | N/A | 0.05221641 | N/A | 0.05509726 | N/A |
| Guaymas Basin Methane G | 330 | N/A | 0.00642214 | N/A | 0.05147287 | N/A | 0.05318532 | N/A |
| Guaymas Basin Methane 4 | 240 | 116 | 0.02751961 | 0.02244724 | 0.05047312 | 0.0951562 | 0.05046968 | 0.17643896 |
| Guaymas Basin Propane | 287 | 124 | 0.00395373 | 0.04127534 | 0.05846108 | 0.08940276 | 0.07136118 | 0.14667714 |
| Guaymas Basin Butane | 260 | 101 | 0.00915535 | 0.02717667 | 0.05765143 | 0.08865665 | 0.07108694 | 0.16991205 |
| Elba Seep | 376 | 202 | 0.02574961 | 0.02232657 | 0.05269071 | 0.09050086 | 0.05543185 | 0.16815102 |
| Menes Caldera 2 | 321 | 204 | 0.011038 | 0.04610654 | 0.04917194 | 0.09313789 | 0.049239 | 0.16557968 |
| Menes Caldera 2A | | 197 | | 0.08124868 | | 0.09037035 | | 0.16776429 |
| Black Sea | 377 | 460 | 0.040185 | 0.02340786 | 0.04503215 | 0.11170009 | 0.03978634 | 0.24550464 |

**Table S3.** Diversity metrics for host and viral communities.

| | Host 16S NMDS, ASV level | Host 16S NMDS, Species level | Viral PCA, Viral genus level | Viral PCA, Viral contig level |
|---|---|---|---|---|
| **Unique energy source** | 0.00039996 | 0.00089991 | 0.9259074 | 0.9429057 |
| **Methane vs non-methane energy source** | 0.00179982 | 0.00189981 | 0.39986 | 0.4427557 |
| **Temperature lifestyle** | 9.999e-05 | 0.00019998 | 9.999e-05 | 9.999e-05 |
| **Original Sampling location** | 9.999e-05 | 9.999e-05 | 9.999e-05 | 9.999e-05 |

**Table S4.** PERMANOVA (ADONIS) values for ordination plots in Figures 2 and S1.

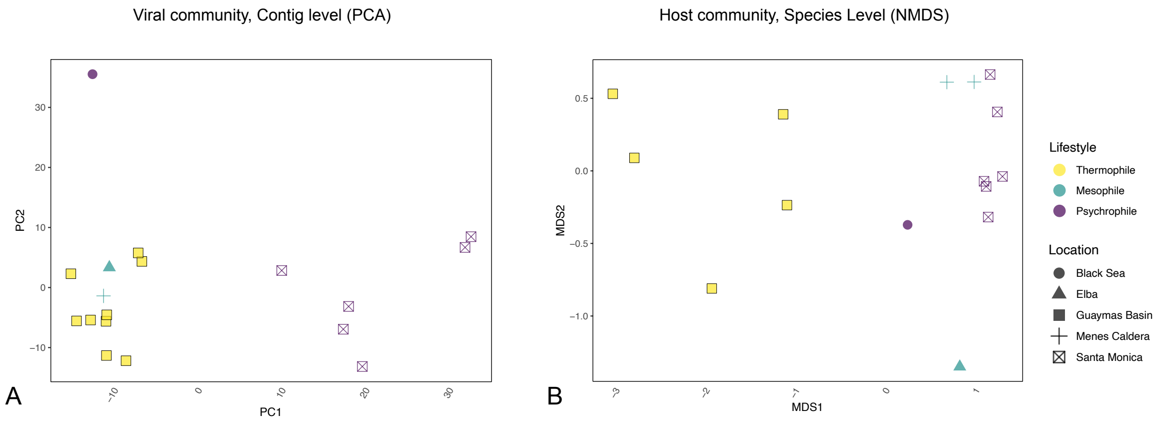


**Figure S1. A:** Viral CLR-transformed abundance-based PCA, at individual viral contig level. PC1 importance: 0.3511, PC2 importance: 0.1547 **B:** 16S relative abundance-based NMDS, at species taxonomy level. Stress=0.076.

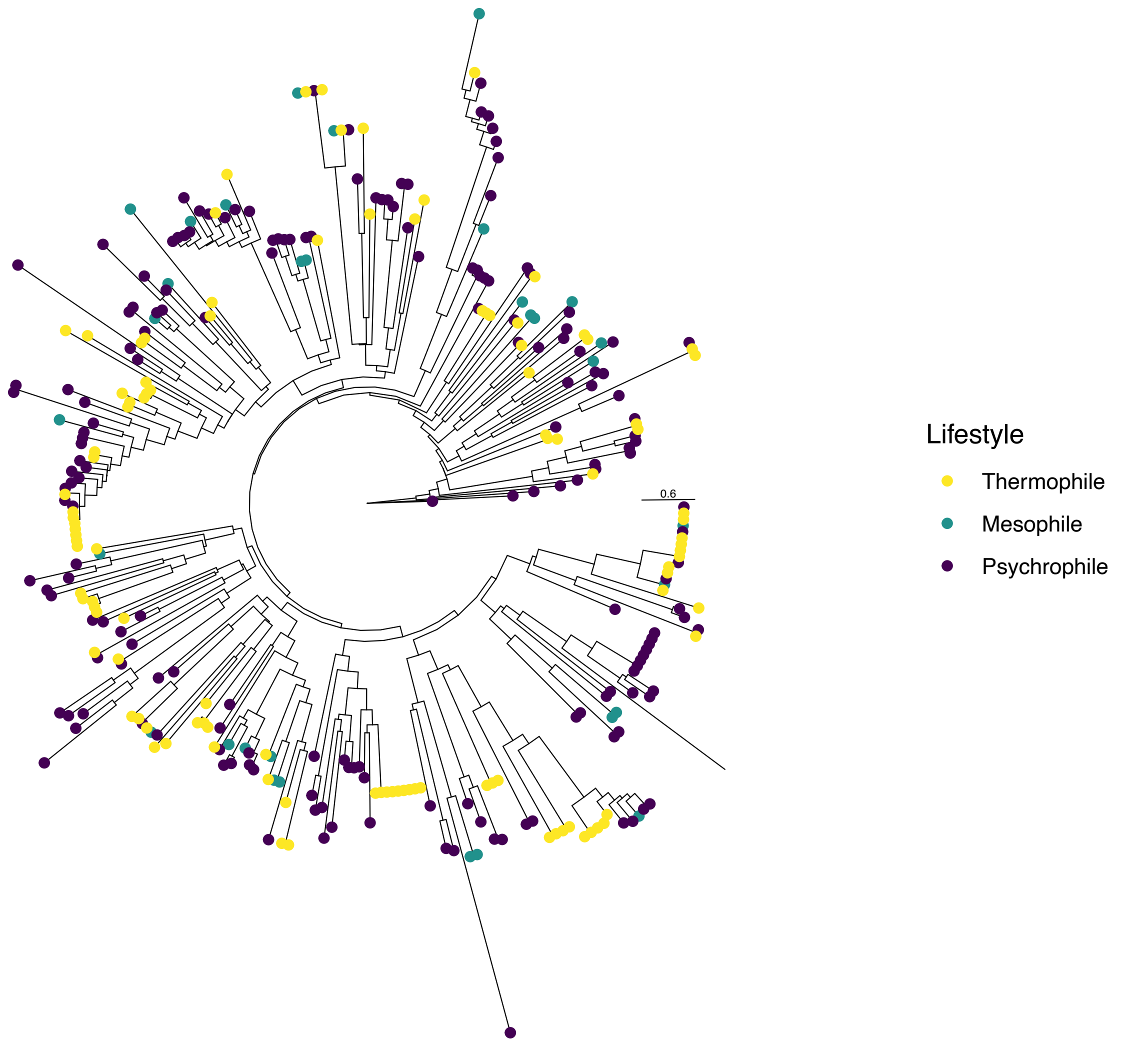


**Figure S2.** Major Capsid Protein Tree, colored by incubation temperature.

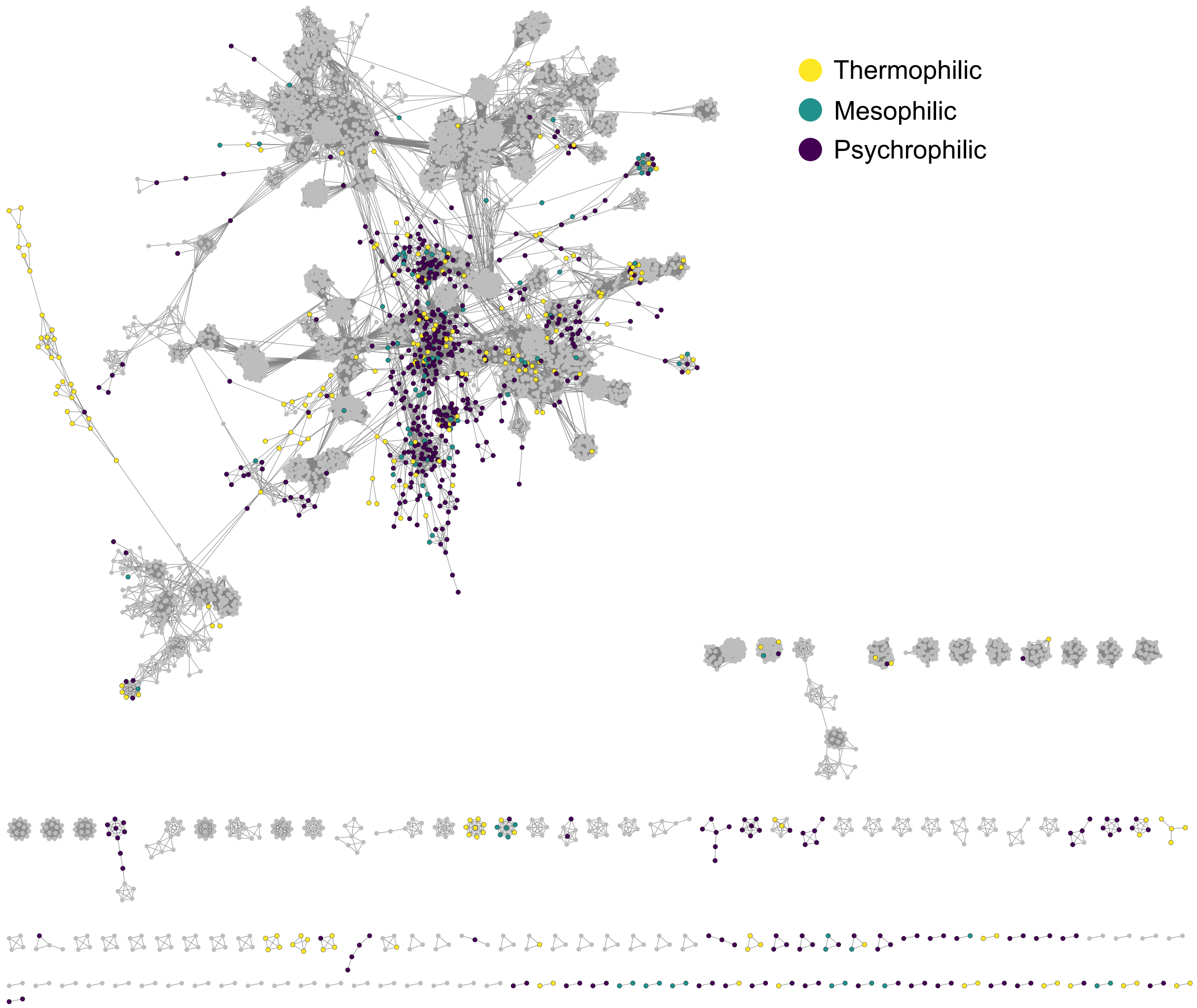


**Figure S3.** Complete vConTACT2 network

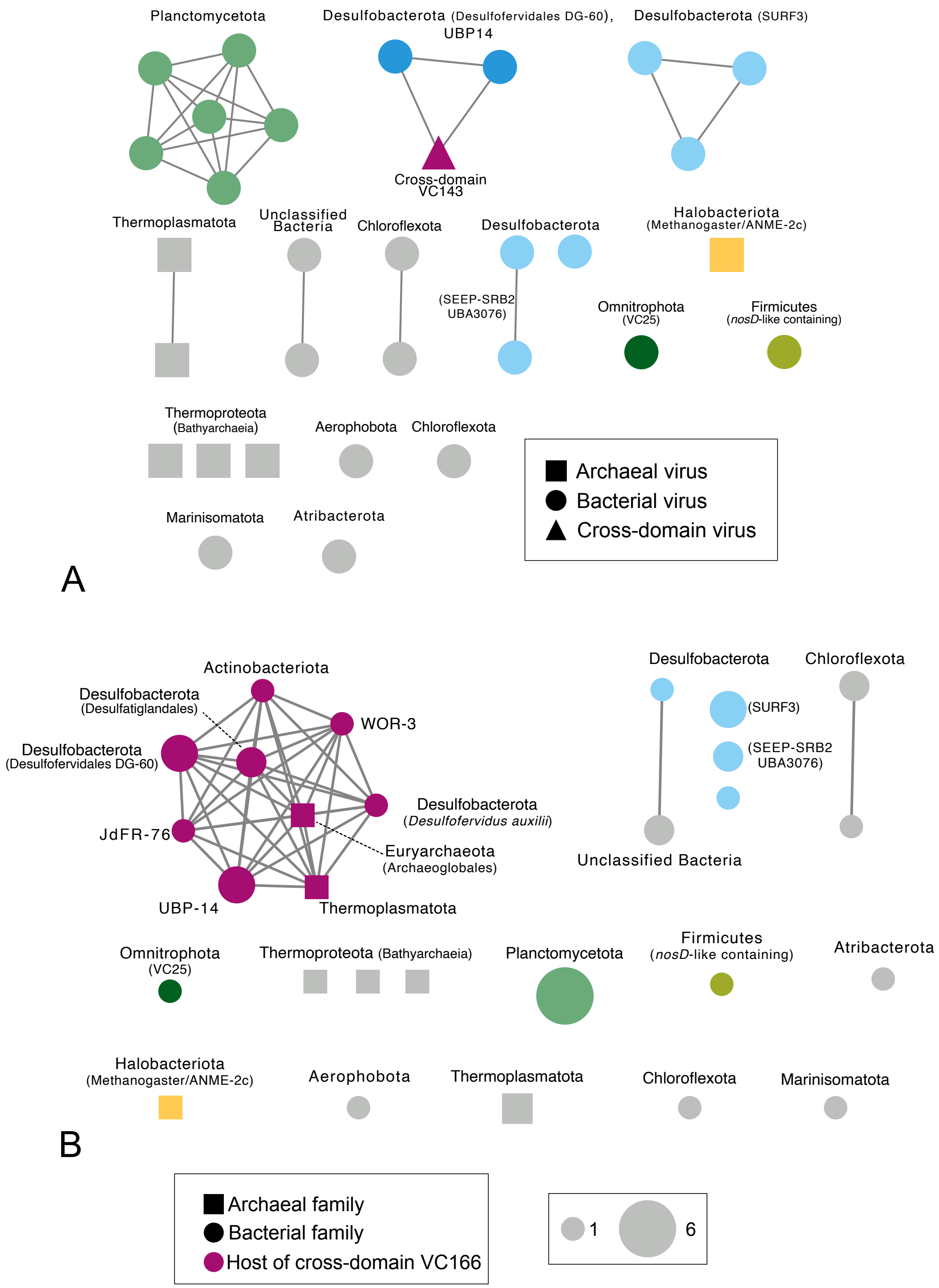


**Figure S4.** Full network of host-phage pairings. **A)** Network with phage genus-level clusters as nodes and shared host at the family level as edges. Nodes are labeled by host phylum; any viral cluster infecting *Desulfobacterota* is colored in shades of blue. **B)** Network with hosts at family taxonomic level as nodes and shared viral clusters as edges. Nodes are labeled by host phylum. Magenta nodes are families that are infected by broad host range viral cluster VC166, though VC166 may not be the only virus infecting that family. Node size corresponds to the number of viral clusters matched to each family via CRISPR spacers or tRNAs.

| ORF | Vibrant Annotation | CD Search (e-value, bitscore) | GTDB r214 BLASTp top phyla (percent identity, e-value, bitscore) | NCBI BLASTp top phyla (percent identity, e-value, bitscore) |
|---|---|---|---|---|
| **45** | K02843 (*waaF*/*rfaF*) | *rfaF* superfamily (COG0859) (1.85e-16, 78.865) | *Desulfobacterota* (43.3, 7.23e-97, 301.0) | *Ca. Methanomethylicota* MAG (47.2, 4e-103, 318), *Thermodesulfobacteriota* MAG (43.31, 2e-97, 304) |
| **57** | K12547 (polysaccharidase *plyA*) | *nosD* (cl34609) with a choice-of-anchor Q domain (cl49486) (1.21e-10, 63.3972) | *KSB1* (39.2, 2.66e-118, 379.0), *Fermentibacterota* (41.7, 8.44e-112, 362.0), *OLB16* (39.5, 1.13e-111, 358.0) | *Bacteroidota* (41.03, 4e-137,428) |
| **74** | K03269 (*lpxH*) | Metallophosphatase superfamily (cl13995 and cl42652) (2.36e-12, 63.5309) | *Thermoplasmatota* (33.2, 1.01e-22, 101.0), *UBP14* (30.8, 1.88e-21, 98.6) | *Bathyarchaeota* MAG (62.26, 3e-93, 283), *Thorarchaeota* MAG (62.26, 1e-92, 182) |
| **82** | K01448 (*amiABC*) | *amiC* family (COG0860) (5.77e-71, 213.204) | *Methanobacteriota* (45.4, 2.43e-37, 140.0, *Bacillota* (45.4, 3.42e-37, 140.0) | *Thermodesulfobacteriota* (*D. auxilii*) MAG (44.89, 5e-38, 140), *Methanobacteriota* (45.65, 1e-37, 143) |
| **112** | Hypothetical protein | Uncharacterized conserved protein w/ vonWillibrand factor type A (cl27002) (9.72e-05, 44.2856) | *Spirochaetota* (e=2.930000e-115, bitscore=360.0) | *Synergistota* MAG (45.66, 2e-140, 426), *Methanobacteriota* MAG (44.06, 2e-139, 423) |
| **109** | K03148 (*thiF*) | E1 activating enzymes of ubiquitin-like proteins (cl22428) (1.84e-15, 71.0409) | *Myxococcota* (34.3, 1.01e-13, 76.3) | *Methanobacteriota* MAG (37.82, 9e-22, 99.0) |
| **16** | K07391 (*comM*) | *YifB* superfamily (cl33973) (5.67e-08, 51.1944) | *Bacillota* (29.0, 3.0e-6, 57.4) | *Deltaproteobacteria* MAG (31.21, 5e-16, 83.2) |
| **84** | K00558 (DNMT1) | *Dcm* DNA-cytosine methylase (COG0270) (7.68e-66, 206.198) | *Halobacteriota* (42.3, 8.71e-50, 173.0) | *Nitrospirota* MAG (41.84, 2e-54, 187) |

**Table S5.** BLAST and conserved domain search results at the phylum level for open reading frames from contig f9bf2df06235b506f7396f56f61038ed in VC143. Though the *plyA* annotation was found to have some conserved domain similarities to *nosD*, it was not found to be a high quality *nosD*-like sequence in a search with the *nosD* HMM. All GTDB results were

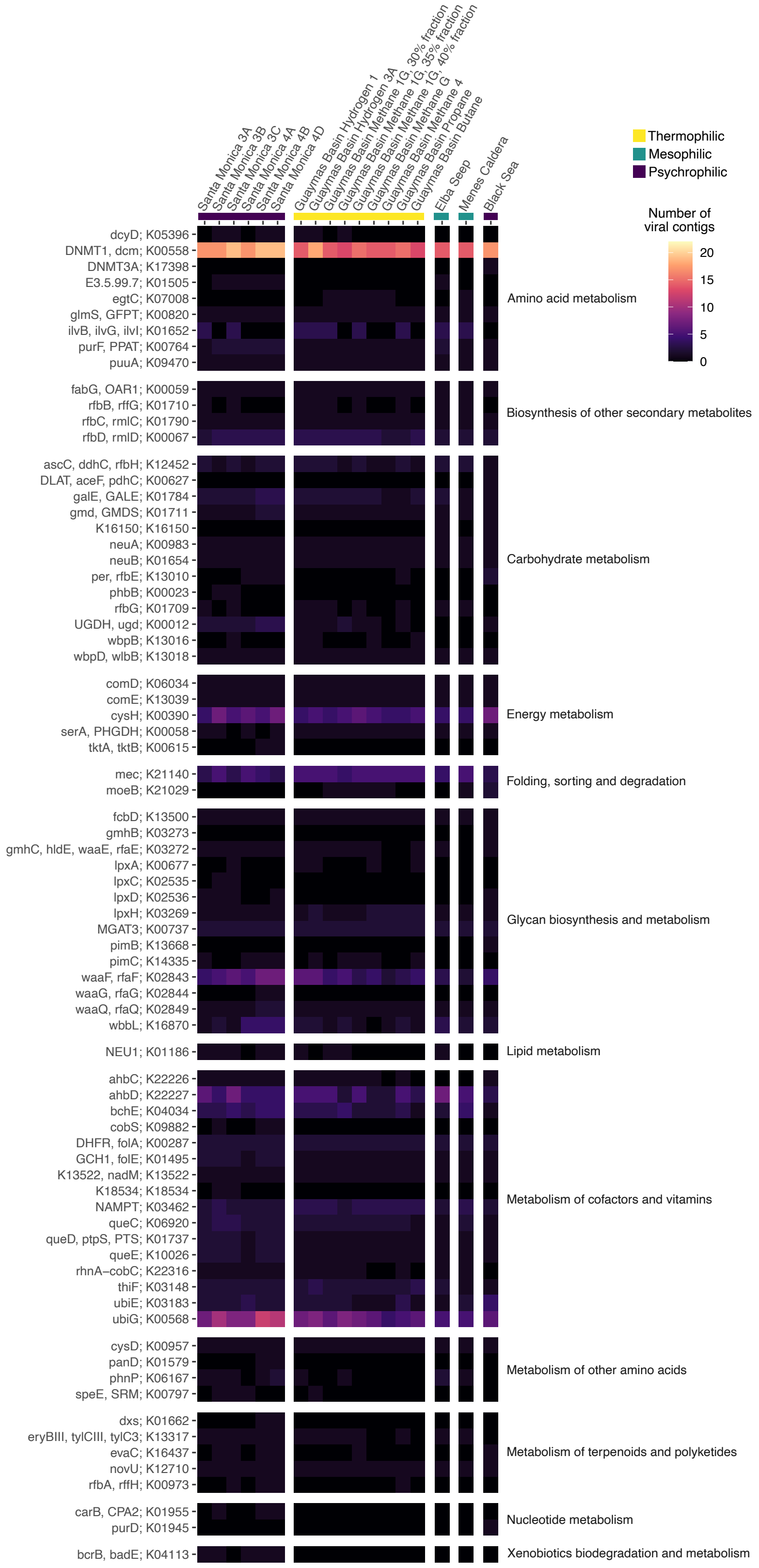


**Figure S5.** Complete list of AMGs and their abundance in each sample. See main text Figure 5 for information on abundance calculations.

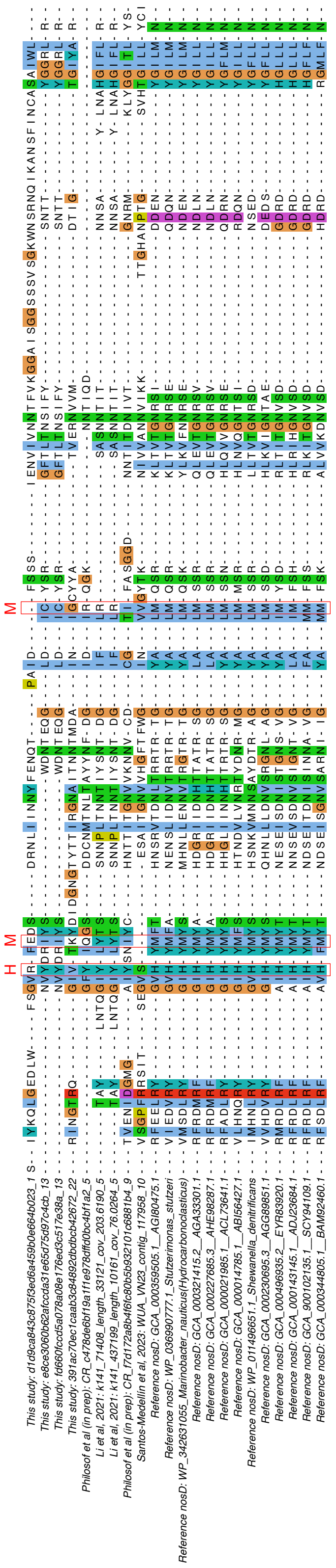


**Figure S6.** Examples of *nosD*-like proteins spanning the conserved His-Met-Met motif. Sequences labeled as *Reference nosD* are canonical *nosD* sequences.

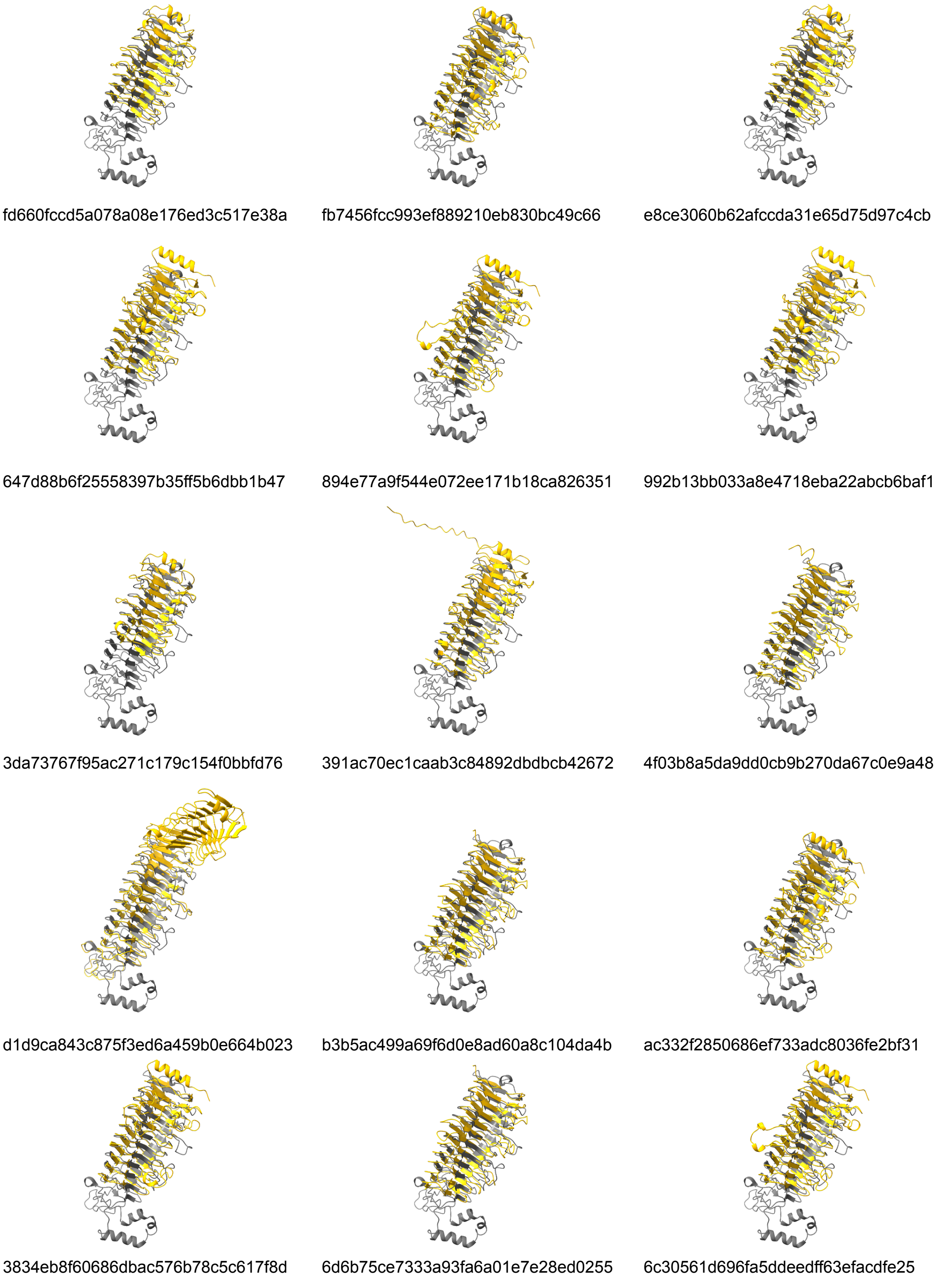


**Figure S7.** Structures of *nosD*-like proteins from this study, labeled by the fasta header of the viral contig from which they came. The *nosD*-like protein is in yellow and is superimposed on the structure of the canonical *nosD* in grey.

### Supplementary Methods

*Development and maintenance of incubations*

Anaerobic alkane-oxidizing enrichments from Guaymas Basin, Elba, Menes Caldera, and the Black Sea were developed via dilution of sediment slurries with anoxic artificial seawater medium: 150mL slurries (150 ml) were transferred in 256 ml serum bottles sealed with butyl rubber stoppers as described in Supplementary Reference #[7]. Enrichments from Santa Monica Basin were developed and maintained as described in Supplementary Reference #[1]. All incubations are maintained in an artificial seawater medium containing sulfate as the sole electron acceptor. Sulfide concentrations were measured to track metabolic activity, and active cultures were consecutively diluted. Spent media from 100mL incubations was replaced by allowing the particles to settle and removing 80mL of medium and replacing the same volume with new anoxic artificial seawater. The *D. auxilii* and *Archaeoglobus* incubations were maintained under 0.2 MPa of 80:20 hydrogen:carbon dioxide, while the AOM enrichments were provided 0.2MPa of 100% methane. Propane and butane enrichments were produced as described above from Guaymas Basin, and slurries and their dilutions were kept at 0.15 MPa propane/butane. A full description is provided in Supplementary Table S1, including references for media composition.

*Media recipe for Guaymas Basin Methane and Hydrogen incubations*

1. Begin with the following in 800mL of water, for a final volume of 1L:

| **Reagent** | **g or mL/L** | **g/mol** | **final mM concentration** |
|---|---|---|---|
| $MgCl_2 \cdot 6H_2O$ | 5.67g | 203.3 | 27.9 |
| $CaCl_2 \cdot 2H_2O$ | 0.22g | 147.0 | 1.5 |
| NaCl | 26.37g | 58.4 | 451.2 |
| KCl | 0.60g | 74.6 | 8.0 |
| $Na_2SO_4$ | 1.50g | 142.0 | 10.6 |
| $K_2HPO_4$ | 0.17g | 174.2 | 1.0 |
| $NH_4Cl$ | 0.11g | 53.5 | 2.0 |
| Se/W solution (see below) | 0.10mL | | 0.01/0.0067 uM |
| Trace elements solution (see below) | 1.00mL | | |
| 250mM HEPES buffer (see below) | 100.00mL | | 25.0 |

2. The pH should be between 7.2-7.45. Filter through a 0.22µm pore size filter to sterilize and flush with nitrogen gas for 20-30 minutes to make the media anoxic.
3. Add the following after the media has been flushed:

| **Reagent** | **g or mL/L** | **final mM concentration** |
|---|---|---|
| 200mM $HS^-$ solution, flushed with nitrogen gas | 2.5mL | 0.5 |
| 1M $NaHCO_3^-$ solution, filter sterilized and flushed with nitrogen gas | 5mL | 5 |

Se/W solution and trace element solution recipes can be found in Supplementary Reference#[8]. For a 500mL stock solution of 250mM HEPES at pH 7.5, use 32.5g HEPES and 2.7g of NaOH.

*Collection and purification of viruses*

Spent media from the enrichment incubations were removed and filtered through a 0.22µm pore size polyethersulfone membrane (Cat. SLGP033N, Millipore Sigma; St. Louis, MO, USA) to remove cells and cell debris. Viruses in the filtrate were concentrated in an Amicon Ultra-15 Centrifugal Filter Unit with a 100kDa cutoff (Cat. UFC910024, Millipore Sigma; St. Louis, MO, USA) spun at 2000×g for 3 minutes (Allegra X-I5R Centrifuge, Beckman Coulter; Brea, CA, USA). To increase yield of viruses from the filtration membrane, the cartridge was wrapped with parafilm and vortexed at the maximum setting for 10 seconds (Vortex Genie 2, Scientific Industries; Bohemia, NY, USA). Concentrated viruses were then purified using ultracentrifugation (Ultima MAX-E Ultracentrifuge; Beckman Coulter, Brea, CA, USA) with Optiprep Density Gradient Medium (Cat. D1556, Sigma Aldrich; St. Louis, MO, USA), according to the protocol adapted from Supplementary Reference #[9] and detailed in Supplementary Reference #[10]. Density fractions were then stained with SYBR™ Gold Nucleic Acid Gel Stain (Cat. S11494, ThermoFisher; Waltham, MA, USA) [11] and visualized with fluorescence microscopy to determine which fractions contained viral particles. The fractions marked by 30%, 35%, and 40% were found to contain the largest number of viral particles. Note that this method does not account for the presence of non-viral particles containing nucleic acid, such as vesicles, but vesicles are thought to affect total viral counts by less than an order of magnitude in seawater samples [12].

*Transmission Electron Microscopy*

Spent media from incubations was collected, filtered, and concentrated as described under *Collection and purification of viruses*. Five microliters of virus concentrate were spotted on a glow-discharged carbon-coated 300 mesh copper TEM grid (Cat. 01813, Ted Pella Inc., Redding, CA, USA). Samples were incubated on the grid at room temperature for five minutes, and excess liquid was wicked off with filter paper. Once dry, grids were stained with 2% uranyl acetate solution in 1X PBS for 30 seconds; the staining procedure was repeated twice for a total of 60 seconds of treatment. Excess stain was removed and grids were air dried completely before imaging with an FEI Tecnai T12 at 120kV. Electron microscopy was performed in the Beckman Institute Resource Center for Transmission Electron Microscopy at Caltech.

*Viral DNA extraction and sequencing*

Density fractions were treated with RNase-free DNaseI (New England Biolabs M0303L) to remove free DNA. Viral DNA was extracted using Promega Wizard PCR Preps DNA Purification Resin (Cat. A7181, Promega, Madison, WI, USA) and Wizard Minicolumns (Cat. A7211, Promega, Madison, WI, USA) in accordance with Supplementary Reference #[13]. Sequencing libraries were prepared using the Illumina DNA Prep library kit (Cat 20018705, Illumina, San Diego, CA, USA) with IDT® for Illumina® DNA/RNA UD Indexes Set C (Cat. 20042666). The DNA from the two Menes Caldera incubations was pooled to ensure sufficient input for library preparation. Paired-end sequencing was performed at the University of Southern California Keck School of Medicine using an S1 flow cell for 300 cycles on the Illumina NovaSeq6000 sequencing platform (v1.5 reagents).

*Cellular DNA extraction and sequencing*

Approximately 15mL of each enrichment was collected and centrifuged at 5000×g and 10˚C for 30 minutes. The supernatant was removed and the resulting cell pellet was extracted using Qiagen's DNeasy Blood and Tissue Kit according to their Gram positive protocol (Cat. #69504, Qiagen, Hilden, Germany). 16S rRNA archaeal and bacterial genes were amplified in duplicate with Q5 Hot Start Master Mix (Cat. M0492S, New England Biolabs, Ipswich, MA, USA) according to the manufacturer's directions, using 515F (5'-TCGTCGGCAGCGTCAGATGTGTATAAGAGACAG-GTGYCAGCMGCCGCGGTAA-3') and 926R (5'-GTCTCGTGGGCTCGGAGATGTGTATAAGAGACAG-CCGYCAATTYMTTTRAGTTT-3') primers [14] that include Illumina adapters for 32 PCR cycles. PCR duplicates were then pooled and barcoded with the Illumina Nextera XT Index Kit v2. Amplification was performed with Q5 Hot Start PCR mixture and 3µL of the pooled duplicate product was added to a 30µL reaction volume, annealed at 66°C, and cycled 11 times. Products were run on a 1.5% agarose gel and quantified by band intensity. These barcoded PCR products were then combined in equimolar amounts into a single tube and 300uL of this pooled sample was run on a 1.5% low melt agarose gel (Cat. BP165-25, ThermoFisher) and purified using Promega's Wizard SV Gel and PCR Clean-up System (Cat. #A928, Promega, Madison, WI, USA). The sample was sequenced by Laragen (Culver City, CA, USA) using the MiSeq Reagent Kit v3 (Cat. #MS-102-3003, 600-cycle) on Illumina's MiSeq platform with the addition of 15-20% PhiX. Figure 1 was generated after dropping singletons and organisms classified as "uncultured" at the family level. The 16S rRNA gene sequencing data was not included for Guaymas Basin Methane 1G and Guaymas Basin Methane G due to a significant temporal gap between virus sampling and 16S rRNA gene sampling, in which the bottles were diluted into larger containers. 16S rRNA amplicon data was processed with the DADA2 v.1.22 workflow [15]. A reproducible workflow is available in the Caltech Library data repository (see Data Availability statement). SILVA SSU database r138 [16] with in-house sequences added was used to assign taxonomy to amplicon sequence variants. Whole genome library preparation and paired-ended sequencing was performed at the Millard and Muriel Jacobs Genetics and

Genomics Laboratory, California Institute of Technology, using a P1 flowcell for 300 cycles with the Illumina NextSeq2000 sequencing platform.

*Cellular metagenome assembly and binning*

Reads were trimmed with bbduk v38.81 [17] and filtered such that reads with <150 nucleotides were eliminated. Reads that passed this step were assembled using metaSPAdes version 3.15.3 [18]. Reads were aligned to the assembled contigs using the Burrows-Wheeler Alignment tool version 0.7.17-r1188 [19] before contigs were binned using MetaBAT version 2.12.1 [20] with a minimum contig length of 1500 bp. The CheckM v1.1.3 lineage workflow [21] was used to determine the quality of each bin. Taxonomy was assigned with GTDB-Tk v2.1.0 [22] using the GTDB R207 database [23–26]. All reads were also mapped back to the full set of contigs using bbmap v38.81 [17]; unmapped reads were reassembled with metaSPAdes.

*vMAG assembly, annotation, and phylogeny*

Viral reads were trimmed using bbduk v38.81 with kmer lengths of 23, minimum kmer length of 11, maximum hamming distance of 1, and parameters set to trim evenly at the overlap of paired reads. Contigs were assembled using metaViralSPAdes v3.15.3 [27] with default parameters. To improve the quality of assembly and number of complete vMAGs, all Guaymas Basin methane incubation reads were pooled before assembly, as were Guaymas Bain hydrogen reads and Santa Monica reads. Viral contigs were identified and quality checked using CheckV [28], viralVerify [27], and seeker [29]. viralVerify was also set to distinguish plasmids. All virome-assembled contigs, regardless of whether they were identified as viral by the preceding algorithms, were subject to annotation by Vibrant [30]; Vibrant filters based on probable viral identity before annotation, adding a fourth set of checks to the three algorithms cited above. All cellular contigs (see *Cellular Metagenome Assembly and Binning*) were checked to identify viruses in the cellular metagenomes; these viral contigs were also annotated with Vibrant and combined with the vMAGs from the viral sequencing for downstream work.

Terminase large subunit (*TerL*) and major capsid protein (*MCP*) phylogenies were constructed using the annotations obtained from Vibrant. Relevant amino acid sequences were aligned using MAFFT v7.505 [31] and trimmed with trimAl v1.4.rev15 [32]. Alignments were manually curated to remove short sequences or sequences with large gaps. Phylogenetic trees were generated using IQ-TREE multicore version 1.6.12 [33].

Auxiliary metabolic genes were included in the analysis if classified as such by Vibrant. Because annotations and open reading frame calls are not always reliable without a great deal of manual curation, AMG abundances per sample were quantified as the abundance of the viral contig on which the AMG was found (see *Viral Abundance Calculations and Diversity*) rather than the abundance of the AMG itself. Within a given sample, a contig was included in the AMG calculations if the centered log ratio of the contig's abundance was >1.

*Viral classification and clustering*

Viral contigs that were 98% complete or higher according to CheckV were classified at the family level using GRAViTy: Genome Relationships Applied to Virus Taxonomy [34], with ICTV Virus Metadata Resource VMR_16-180521_MSL36 (https://ictv.global/vmr, [35]). The whole-genome protein clustering network was generated using vConTACT2 [36] using Refseq release 207, pcs mode MCP, vcs mod ClusterONE. The network was visualized using Cytoscape. ANME-1 phages from Supplementary Reference#[37] were included in the network calculation due to the similarities of the sampling site and host taxonomy to those used in this study. T7 control sequences were included in vConTACT2 analysis as an internal check.

*Viral abundance calculations and ordination*

Viral reads were quantified per viral cluster and per contig using salmon v1.10.0 [38] and summarized using tximport v1.18.0 [39]. DESeq2 v1.30.1 [40] was used to quantify the abundance of each cluster/contig per library. Abundances were centered log ratio (CLR) transformed with ALDEx2 v1.25.1 [41]. Diversity metrics were calculated using the transcripts per million value for each viral cluster, as output by tximport. Principal components, corresponding PERMANOVA (Adonis, analysis of dissimilarities) values, and rank abundances were calculated using the CLR-transformed data.

Host community nonmetric multidimensional scaling (NMDS) ordination plots were generated using the vegan v2.6-4 [42] metaMDS function (Bray-Curtis dissimilarity metric on relative abundance data) at the amplicon sequence variant (ASV) level [43]. Viral community principal component analysis (PCA) ordination plots were generated using the prcomp function (Euclidean distance on centered log ratio transformed abundance data) at the viral genus level; principal component analysis (PCA) was used for the viral community instead of NMDS because the centered log ratio transformation moves compositional data into Euclidean space [44]. For both host and virus communities, corresponding PERMANOVA values were calculated with the vegan adonis2 function.

*CRISPR and tRNA identification and host matching*

Potential cellular host genomes were obtained from NCBI from Projects PRJNA276404 [2, 45]; PRJNA318983 & PRJNA319143 [4]; PRJNA418316 [46]; PRJEB36446 & PRJEB36096 [47]; PRJNA713414 [48]; PRJNA758896 [1]; PRJNA875076 & PRJNA721962 [37]; members of *Ca. Desulfofervidaceae* from PRJNA762493 [49]; and personal communications with Dr. Daan Speth at the University of Vienna and Dr. Rafael Laso-Pérez at the Museo Nacional de Ciencias Naturales, Madrid. These genomes were generated from bottles used in this study or were sampled from closely-related sites. Sixty-three additional *Archaeoglobus* genomes were obtained from GTDB under the search time “archaeoglobus”. Additional cellular genomes were generated from three of the incubation bottles (Guaymas Basin Methane 1G, Guaymas Basin Methane G, and Guaymas Basin Hydrogen 3A) as detailed above.

CRISPRs were called using CRISPRDetect 3.0 [50] and CRISPRCasTyper v1.1.4 [51]. Host genome CRISPR spacers were combined and dereplicated with MMseqs2 [52] version 13.45111easy-cluster workflow at 99% minimum sequence identity and 80% coverage. BLASTn v2.12.0+ and SpacePharer v5.c2e680a [53] were used to compare the viral contigs and the host spacers. BLAST hits were considered high-quality with at least 98% sequence similarity and no more than 2 mismatches.

tRNAs were identified in putative host genomes and viral genomes using tRNAscan-SE v2.0.5 [54] and Aragorn v1.2.41 [55]. Viral and host tRNAs were combined and dereplicated respectively, as done for the CRISPRs. Host tRNAs were aligned against viral contigs and viral tRNAs using BLAST. Matches were considered high-quality only if they were exact matches: 100% sequence identity, 100% coverage, and no mismatches.

Host/phage network diagrams were made using adjacency matrices of viral clusters that shared hosts, and hosts infected by the same viral clusters, respectively. Where viruses were not clustered at the genus level by vConTACT2, the contig was used on its own. Networks were calculated using the networkX from_pandas_adjacency function [56], and self-loops were removed. The network was visualized using Cytoscape [57]. The genome map of VC143 was visualized using DNA Features Viewer [58].

*Identification of nosD-like proteins using a hidden Markov model*

Input sequences for the *nosD* Hidden Markov Model were derived from Supplementary Reference #[59] Extended Data Figure 7A. Sequences for *Geobacillus thernodenitrificans* (WP_008880109.1), *Shewanella denitrificans* (WP_011496651.1), *Stutzerimonas stutzeri* (WP_036990777.1), *Marinobacter nauticus* (WP_342631055.1), *Wolinella succinogens* (WP_011138819.1), *Hydrogenobacter thermophilus* (WP_012962810.1), and *Deslfitobacterium* hafniense (WP_018214548.1) were aligned against a curated set of genomes (See Caltech Library link under Data Availability) using BLAST v2.9.0+. Hits with e-value<0.0005 and bitscore>200 were retained and aligned with the selected sequences from Supplementary Reference #[59] using MUSCLE v3.8.31 [60]. The alignment was manually curated and dereplicated at 70% identity in Jalview [61].

The HMM was built from this alignment; all viral proteins output by Vibrant were searched for *nosD*-like proteins [62]. *nosD*-like proteins were included in this study if they had an hmmsearch e-value<0.001, came from contigs that were confirmed to be viruses using the criteria under *vMAG Assembly and Annotation*, came from contigs at least 5000 basepairs in length, and had a greater proportion of viral genes than host genes according to checkV. Vibrant *nosD* annotations that were not found in the hmmsearch results were included in the study if they were >200 amino acids in length and appeared to span the conserved His-Met-Met residues of the canonical *nosD* proteins in a MUSCLE-generated alignment. These open reading frames were truncated to remove any additional protein domains before inclusion in the phylogeny; the presence of additional domains and the location within the ORF of the nosD-like protein were determined using NCBI's Conserved Domain Search.

Viral *nosD*-like proteins from the Tara Oceans and Malaspina viral study [63], Costa Rica methane cold seeps [64, 65], lake sediment and water column [66], and soils [67] were similarly searched using the *nosD* HMM and thresholded at e-value<0.001. For published datasets that were only available as nucleic acid fasta files, genes were predicted with Prodigal V2.6.3: February, 2016 [68] before hmmsearch was run. Alignments were generated with MUSCLE v3.8.31 [60] and trees were inferred using IQ-TREE2 [33] using the model finder mode and confidence in tree topology was evaluated with an ultrafast bootstrap value of 1000. Protein structures were predicted using Alphafold2 [69] in monomer mode with the reduced set of databases (MGnify, PDB70 (with PDBx/mmCIF and PDB SEQRES), BFD, Uniclust30, Uniprot, and Uniref90) and visualized using ChimeraX [70–72].
